# *Plasmodium falciparum* infection during pregnancy impairs fetal head growth: prospective and populational-based retrospective studies

**DOI:** 10.1101/203059

**Authors:** Jamille Gregório Dombrowski, Rodrigo Medeiros de Souza, Flávia Afonso Lima, Carla Letícia Bandeira, Oscar Murillo, Douglas de Sousa Costa, Erika Paula Machado Peixoto, Marielton dos Passos Cunha, Paolo Marinho de Andrade Zanotto, Estela Bevilacqua, Marcos Augusto Grigolin Grisotto, Antonio Carlos Pedroso de Lima, Julio da Motta Singer, Susana Campino, Taane Gregory Clark, Sabrina Epiphanio, Lígia Antunes Gonçalves, Cláudio Romero Farias Marinho

## Abstract

**Background:** Malaria in pregnancy is associated with adverse effects on the fetus and newborns. However, the outcome on a newborn’s head circumference (HC) is still unclear. Here, we show the relation of malaria during pregnancy with fetal head growth.

**Methods:** Clinical and anthropometric data were collected from babies in two cohort studies of malaria-infected and non-infected pregnant women, in the Brazilian Amazon. One enrolled prospectively (PCS, Jan. 2013 to April 2015) through volunteer sampling, and followed until delivery, 600 malaria-infected and non-infected pregnant women. The other assembled retrospectively (RCS, Jan. 2012 to Dec. 2013) clinical and malaria data from 4697 pregnant women selected through population-based sampling. The effects of malaria during pregnancy in the newborns were assessed using a multivariate logistic regression. According with World Health Organization guidelines babies were classified in small head (HC < 1 SD below the median) and microcephaly (HC < 2 SD below the median) using international HC standards.

**Results:** Analysis of 251 (PCS) and 232 (RCS) malaria-infected, and 158 (PCS) and 3650 (RCS) non-infected women with clinical data and anthropometric measures of their babies was performed. Among the newborns, 70 (17.1%) in the PCS and 934 (24.1%) in the RCS presented with a small head (SH). Of these, 15 (3.7%) and 161 (4.2%), respectively, showed microcephaly (MC). The prevalence of newborns with a SH (30.7% in PCS and 36.6% in RCS) and MC (8.1% in PCS and 7.3% in RCS) was higher among babies born from women infected with *Plasmodium falciparum* during pregnancy. Multivariate logistic regression analyses revealed that *P. falciparum* infection during pregnancy represents a significant increased odds for the occurrence of a SH in newborns (PCS: OR 3.15, 95% CI 1.52-6.53, p=0.002; RCS: OR 1.91, 95% CI 1.21-3.04, p=0.006). Similarly, there is an increased odds of MC in babies born from mothers that were *P. falciparum-infected* (PCS: OR 5.09, 95% CI 1.12-23.17, p=0.035). Moreover, characterization of placental pathology corroborates the association analysis, particularly through the occurrence of more syncytial nuclear aggregates and inflammatory infiltrates in placentas from babies with the reduced head circumference.

**Conclusions:** This work indicates that falciparum-malaria during pregnancy presents an increased likelihood of occurring reduction of head circumference in newborns, which is associated with placental malaria.

**Trial Registration:** registered as RBR-3yrqfq in the Brazilian Clinical Trials Registry

## BACKGROUND

Malaria remains a major global health problem, with approximately one billion people living at high-risk of being infected (World Health Organization. 2017). *Plasmodium* spp. infection impacts the health of the poorest and marginalized communities in the endemic countries, particularly in infants and pregnant women, with around 125 million pregnancies at risk of infection each year (Dellicour et al. 2010). Malaria during pregnancy, especially falciparum-malaria, can be devastating and fulminant, leading to high mortality for both mother and fetus (Desai et al. 2007). During pregnancy, the infected erythrocytes accumulate and sequester in the placental intervillous space, causing placental histopathological changes, which triggers an exacerbated inflammatory response that is highly detrimental (Ismail et al. 2000). The deleterious effects caused by malaria infection during pregnancy depend on various factors, such as the woman’s immunity, the number of previous pregnancies and the trimester of gestation, with primigravida and secundigravida most susceptible and suffering the greatest consequences (Rogerson et al. 2007).

A heightened inflammatory response perturbs the maternal-fetal interface and impairs critical placental functions. Therefore, maternal malaria presents a major impact on fetus and newborns, being the main cause of abortion, stillbirth, premature delivery and fetal death in malaria-endemic countries (Desai et al. 2007). Low birth weight (LBW) caused by prematurity or intrauterine growth retardation (IUGR) is commonly observed in babies born from mothers who had malaria during pregnancy, contributing to around 100,000 infant deaths each year (Guyatt and Snow 2001; Desai et al. 2007; Rogerson et al. 2007). Additionally, *in utero* exposure to malaria parasites has been shown to impact the fetus or newborn head circumference (HC), a proportional reduction as an outcome of the IUGR (Menendez et al. 2000; Meuris et al. 1993). Albeit, no further studies have tried to unpick a specific disproportionate HC reduction associated with malaria during pregnancy.

Several studies have reported the association of intrauterine infections with a high risk of the newborn to have LBW and brain injury (Zhao et al. 2013). A group of microorganisms designated as TORCH, an abbreviation for *Toxoplasma*, rubella, cytomegalovirus, and Herpes simplex that now also comprise *T. pallidum* (Syphilis), hepatitis virus, and HIV, and recently, the Zika virus are frequently associated with reduced HC in newborns (Neu, Duchon, and Zachariah 2015; Tetro 2016). The more adverse consequence that results from these infections is microcephaly at birth, which is defined by a reduction of the occipitofrontal HC of more than two standard deviations (SD) below the median compared to age and sex-matched control population (Passemard, Kaindl, and Verloes 2013). Although the brain insult is defined by the cranium size, it also reflects a reduction of the brain volume and an impairment of cognitive abilities (Passemard, Kaindl, and Verloes 2013).

Thus, to investigate the relation of malaria during pregnancy on the fetus head growth, we analyzed data from a prospective and a retrospective cohort from newborns delivered between 2012 and 2015 in Cruzeiro do Sul (Acre State in the Southwestern Brazilian Amazon Basin), where 46% of the total falciparum-malaria Brazilian cases occur (SIVEP - Secretaria de Vigilância em Saúde - Ministério da Saúde 2015; Ferreira and Castro 2016).

## METHODS

### Setting

Two cohort studies were conducted in the Amazonian region of the “Alto do Juruá” valley (Acre, Brazil), evaluating maternal-child pairs data of births at the general maternity ward, Hospital da Mulher e da Criança do Juruá (HMCJ, Cruzeiro do Sul), where approximately 90% of the total deliveries in the region occur. “Alto do Juruá” valley is in the extreme southwest of the Brazilian Amazon Basin, covering an area of 74,965 km^2^, predominantly rainforest, and a population of ~200,000 inhabitants. It is limited to the north by the Amazonas state, to the east by the Acre Valley (Acre), and to the south and west by Peru (Fig. 1). This is a region of high malaria endemicity in Brazil, with an annual parasite incidence above 100, where *P. vivax* is responsible for 70-80% of the malaria cases, and where 46% of the total *P. falciparum* Brazilian cases occur (Ferreira and Castro 2016; Kohara Melchior and Chiaravalloti Neto 2016). In this region, 18% of women acquire *Plasmodium* infection during pregnancy (SIVEP - Secretaria de Vigilância em Saúde - Ministério da Saúde 2015).

**Figure 1.**
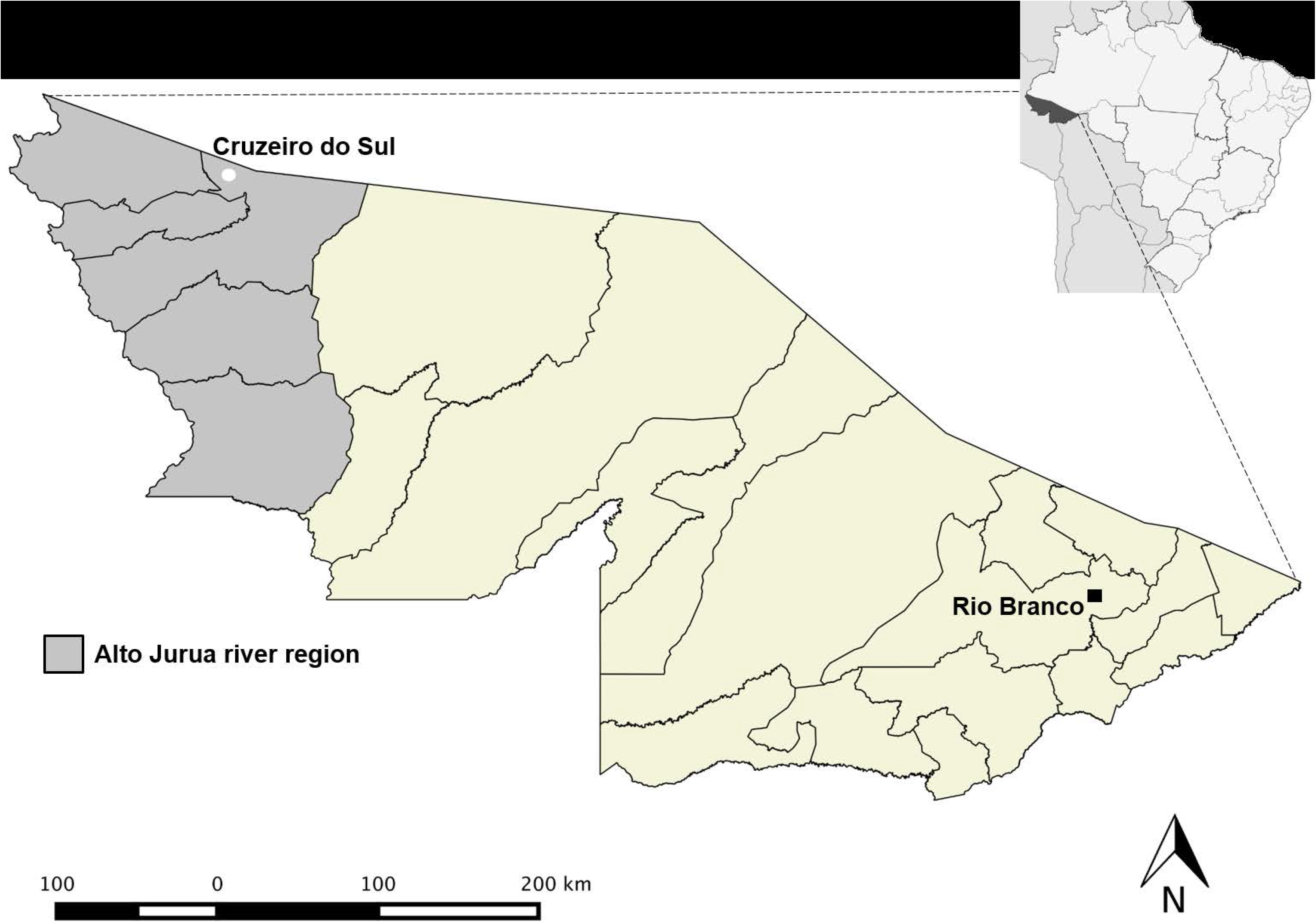
Map showing the location of the field site, Alto do Juruá river region, Northwest of the Acre State, Brazilian Amazon. The map also indicates Cruzeiro do Sul where the field laboratory is situated, and Rio Branco, the capital of the state of Acre.

### Prospective cohort study (PCS)

#### ▪ Design and participants

A total of 600 pregnant women were enrolled through volunteer sampling of equal numbers of *P. falciparum-, P. vivax*-infected, and non-infected pregnant women, and followed until delivery, between January 2013 and April 2015. The women were recruited during their first pregnancy visit to the antenatal care (ANC) clinic. Each pregnant woman was followed by a trained nurse, which involved at least two domiciliary visits, at the second and third trimester, to monitor their clinical state, in addition to the usual prenatal care in health care services.

#### ▪ Samples collection

At the time of recruitment, data was collected on socioeconomic, clinical, and obstetric variables, and peripheral blood and thick and thin blood smears were used to diagnose and confirm malaria infection. During the domiciliary visits, clinical and obstetric data were obtained, and collected a peripheral blood sample. An additional blood sample was collected in each episode of malaria during pregnancy. At the time of delivery, clinical data were collected from mother and newborn, as well as a placental biopsy and blood samples.

#### ▪ Samples processing

The peripheral and placental blood was collected in heparin tubes and then separated into plasma and whole blood cells using a centrifuge. Thin and thick blood smears were stained with Giemsa. The placental biopsies were fixed in 10% neutral buffered formalin at 4°C until they could be sent to the University of São Paulo for processing. Paraffin-embedded 5μm sections of placental tissue were stained with Hematoxylin-Eosin (H&E) or Giemsa for histological examination. Total DNA was obtained from whole blood cells using a commercially available extraction kit (QIAmp DNA Mini Kit, Qiagen), following the manufacturer’s instructions.

#### ▪ Gestational age estimation

The gestational age of all women from the PCS was estimated by woman’s last menstrual period (LMP) and adjusted by ultrasound during the first trimester of pregnancy.

#### ▪ Newborns classification according to head circumference

Based on the gestational age, and on the HC size and gender, each newborn from the PCS was assigned into groups using the INTERGROWTH-21^st^ Project (Villar et al. 2014). An individual was in a normal head circumference (NHC) range if their HC was within one SD of the median. Newborns with HC below one SD below the median were considered to have a small head (SH) (Brennan, Funk, and Frorhingham 1985). Newborns with HC below two SD below the median were classified as having microcephaly (MC) (Passemard, Kaindl, and Verloes 2013).

#### ▪ Screening of malaria infection

Malaria during pregnancy was diagnosed from thin and thick blood smears by two experts in microscopy of the endemic surveillance team of Cruzeiro do Sul (Acre, Brazil). Furthermore, all samples collected throughout the pregnancy were screened for the presence of malaria parasites, by microscopy and confirmed by a real-time PCR technique (PET-PCR). This technique detects in multiplex the *Plasmodium* spp. and *P. falciparum*, and in singleplex *P. vivax* if only *Plasmodium* spp. is detected in the first PCR. PET-PCR has a detection limit of 3.2 parasites/ml (Lucchi et al. 2013). The real-time PCR was performed on the 7500 Fast Real-Time PCR System (Applied Biosystems, ThermoFisher). All the women who had malaria during pregnancy were treated with antimalarial drugs under medical prescription, according to the Brazilian Ministry of Health (MoH) guidelines, with further treatment confirmation.

#### ▪ Histopathology evaluation

The histopathologic examination involved using placental tissue slides. The Hematoxylin-Eosin-staining allowed evaluating the syncytial nuclear aggregates (SNA), fibrinoid necrosis, and fibrin deposition (Souza et al. 2013). The hemozoin presence was assessed through microscopy of polarized light (Romagosa et al. 2004). The leukocyte (CD45) and monocyte inflammatory infiltrate (CD68), and the villous vascularity (CD31) have been evaluated by immunohistochemistry using the tissue microarray (TMA) technique, conducted at the AC Camargo Hospital, in São Paulo, Brazil, as described elsewhere (Ataíde et al. 2015; Hsu, Raine, and Fanger 1981). The proliferation index was calculated through quantitative image analysis of anti-Ki-67/DAB staining (Tuominen et al. 2010). Additional file 1 describes these procedures in detail. The images of placenta were captured by a Zeiss Axio Imager M2 light microscope equipped with a Zeiss Axio Cam HRc camera and analyzed by Image J software (http://imagej.nih.gov/ij).

#### ▪ Angiogenic factors and Leptin measurement

The angiogenic factors, vascular endothelial growth factor A (VEGFA, and it receptors VEGFR1/FLT1 and VEGFR2/FLK1), angiopoietins 1 and 2 (ANG-1 and ANG-2, and their associated soluble receptor the TEK receptor tyrosine kinase (TIE-2)), and the leptin hormone were measured in placental plasma (1:20 dilution for all factors) using the DuoSet ELISA development kits (R&D), according to manufacturer’s guidelines.

#### ▪ Screening of other infectious agents

All pregnant women were screened in the local ANC clinics for toxoplasmosis, hepatitis, syphilis, and HIV by measuring antibodies titers, following the Brazilian MoH guidelines. Further, peripheral plasma from women that delivered babies with small head and microcephaly, irrespective of the infection status and *Plasmodium* species, was tested to confirm the absence of other infectious agents during pregnancy. Tests for *Toxoplasma gondii*, Rubella, Cytomegalovirus, Herpes simplex virus, Syphilis, HIV, Dengue virus, Chikungunya virus, and Zika virus were performed retrospectively by ELISA assays in peripheral blood collected until the 28 weeks of gestation. In pregnant women that delivered babies with microcephaly, plasma samples of two different time points of the pregnancy were tested. All the serological tests were performed using commercially available kits: HIV 1/2 and total Syphilis (Symbiosys) and IgG/IgM to Toxoplasmosis, Rubella, Cytomegalovirus and Herpes simplex (TORCH) (Virion\Serion), and used according to the manufacturer’s instructions. To detect Dengue, Chikungunya, and Zika current viral infections, qualitative assays were carried out by IgM capture using a specific viral antigen for DENV, ZIKV, and CHIKV, as previously described (Sow et al. 2016). The identification of specific IgG antibodies to CHIKV was performed using a specific viral antigen (Sow et al. 2016), and to DENV and ZIKV were made with an antigen derived from a whole DENV-2 NS1 protein and a portion of the NS1 protein, respectively (unpublished data). Developing color was quantified on an automatic microliter plate reader Spectramax Plus 384 (Molecular Devices). The results were expressed as optical density (OD) at 405/630 nm or 450/630 nm (Virion/Serion and Symbiosys/Alka Kits, respectively). In TORCH analyses, the presence of IgG and IgM antibodies were classified as positive, negative or borderline according to an OD range adopted by standard positive control mean. For Rubella and *Toxoplasma gondii* (IgG) avidity test was performed according to the manufacturer’s specifications (Virion/Serion), and in all TORCH IgM tests, we use the rheumatoid factor absorbent reagent (# Z200, Virion/Serion). All the kits followed the validation criteria, and the presence of IgG and IgM antibodies for Syphilis and HIV antigens were determined by comparing the absorbance value of serum samples with the cut-off value of standards of reference controls and classified as positive or negative. All tests were performed without the operator knowledge of the group classification for each sample. If the test was inconclusive the screen was repeated using samples from two different gestational time-points. Newborns were excluded from the analysis whenever their mothers presented antibody titers for IgM.

#### ▪ Measurement of cytokines/anaphylatoxins by bead array

The levels of the cytokines IL-12p70, TNF, IL-10, IL-6, IL-1b, and IL-8 in the placental plasma, were detected and quantified by a CBA human inflammatory kit (BD Biosciences) that was used according to the manufacturer’s protocol. For complement activation studies (measuring C3a, C4a, and C5a) the CBA human anaphylatoxin kit (BD Biosciences) was used. The samples were analyzed in a two-laser BD FACSCalibur flow cytometer with CellQuest version 5.2 software (BD Biosciences), and concentrations computed using FCAP array software version 3.0.1 (BD Biosciences). All plasma samples were processed and kept at -80°C in Cruzeiro do Sul until they were sent to the University of São Paulo.

### Retrospective cohort study (RCS)

#### ▪ Design, participants and data collection

A total of 4697 maternal-child pairs were selected retrospectively through a population-based sampling of all deliveries occurring between January 2012 and December 2013. The data from the Brazilian Epidemiological Surveillance Information System (SIVEP)-Malaria of the mother malaria infection status during pregnancy was assembled with the clinical and anthropometric data present in the medical records of the mother and the newborn. This was followed by the collection and collation of the data to evaluate the newborns further.

#### ▪ Gestational age estimation

The gestational age in the RCS was established by the woman’s last menstrual period (LMP). These data were obtained from the medical records. The LMP method is recommended by the Brazilian MoH for gestational age calculation when it is not possible to use ultrasound.

#### ▪ Newborns classification according to head circumference

Based on the gestational age estimation methodologies, and on the HC size and gender, each newborn from RCS was assigned into groups using the WHO child growth standards (WHO-CGS) (WHO Multicentre Growth Reference Study Group 2007). Gestational age assessment is considered accurate when acquired through ultrasound performed early in the first trimester, but the date of the last menstrual period is considered unreliable (World Health Organization 2016). According to WHO guidelines, the WHO-CGS provides an appropriate reference standard for term neonates when gestational age is not reliably known. An individual was in a normal head circumference (NHC) range if their HC was within one SD of the median, (boys 33.2 ≥ HC ≤ 35.7, girls 32.7 ≥ HC ≤ 35.1). Newborns with HC below one SD below the median were considered to have a small head (SH) (boys HC < 33.2, girls HC < 32.7) (Brennan, Funk, and Frorhingham 1985). Newborns with HC below two SD below the median were classified as having microcephaly (MC) (boys HC < 31.9, girls HC < 31.5) (Passemard, Kaindl, and Verloes 2013).

#### ▪ Screening of malaria infection

Malaria during pregnancy was diagnosed from thin and thick blood smears by microscopists of the endemic surveillance team of Cruzeiro do Sul (Acre, Brazil), whenever women show suspicious malaria symptoms. These data were obtained from the Brazilian Epidemiological Surveillance Information System (SIVEP)-Malaria. All the women who had malaria during pregnancy were treated with antimalarial drugs under medical prescription, according to the Brazilian MoH guidelines.

#### ▪ Screening of other infectious agents

All pregnant women were screened in the local ANC clinics for toxoplasmosis, hepatitis, syphilis, and HIV by measuring antibodies titers, following the Brazilian MoH guidelines.

### Newborn anthropometric measures

In the two cohort studies, PCS and RCS, the newborn anthropometric measures were obtained immediately after the delivery, maximum within 24h, by trained nurses. Weight was measured in grams (g) using digital pediatric scales, with a precision of 5 g, and the length and occipitofrontal head circumference (HC) were measured in centimeters (cm), using a non-stretching flexible measuring tape. Rohrer’s ponderal index is the newborns’ weight in grams divided by the cube of the length in centimeters, and babies are considered proportional when values are above 2.5, corresponding to the 10^th^ percentile (WHO Expert Committee on Physical Status 1995). An Apgar score indicates the physical condition of the newborn, relative to its response to stimulation, skin coloration, heart rate, respiratory effort, and muscle tone. If the Apgar Score is between 7 and 10 the newborn is considered normal; if it is between 4 and 6 it is indicative that some assistance for breathing might be required; and below 4, the baby needs several interventions (American Academy of Pediatrics Committee on Fetus and Newborn and American College of Obstetricians and Gynecologists Committee on Obstetric Practice 2015).

#### Exclusion criteria

Our analysis was restricted to babies that had been born at term (37 – 42 weeks of gestation) with at least 2500 grams of weight in a single birth and from mothers of fertile age (13 – 47 years old). Women were excluded if they had a history during pregnancy of smoking, drug use and/or alcohol consumption, and who presented with infections (TORCH, HIV, Hepatitis B virus, Hepatitis C virus, Syphilis, Dengue, Chikungunya and Zika virus), and/or other comorbidities (e.g. hypertension, pre-eclampsia/eclampsia, diabetes mellitus, preterm delivery, stillbirth, and newborn with congenital malformation). Due to the extremely high percentage of C-sections performed in Brazilian maternity units, women who underwent a C-section were not excluded from the study.

#### Statistical analyses

Data were analyzed using R (r-project.org), Stata (StataCorp), Minitab 18 and GraphPad Prism software. Continuous variables were summarized using means and SD, medians, and interquartile ranges (IQR). Categorical variables were summarized using frequencies and percentages. Differences between groups were evaluated using Mann-Whitney U-tests accordingly. Categorical data and proportions were analyzed using chi-square tests. All *p*-Values were 2-sided, at a significance level of 0.05. To assess the association between malaria and microcephaly, adjusted odds ratios (OR) with 95% confidence intervals (CI) were estimated using a multivariate logistic regression approach. These models included infection by malaria (no/yes), maternal age (≥ 18 years old / ≤ 17 years old) and the number of gestations (two or more/one) as explanatory variables and SH (yes/no) or microcephaly (yes/no) as response variables. The first category for each explanatory variable was considered as reference (Hosmer and Lemeshow 2013). Missing data were imputed or “filled in” within a multiple imputation framework using the “MICE” library within the R software (Rubin 1996; Van Buuren and Groothuis-Oudshoorn 2011). In particular, 5 datasets were completed and the results pooled across allowing for the uncertainty in the imputation process.

The current sample sizes present a deviation from those proposed at the outset. It was proposed to enroll ~400 infected and ~800 non-infected pregnant women into the prospective cohort study. We were unable to recruit to this 2:1 ratio, as some initially included in the non-infected group, were transferred to an infected group upon *Plasmodium* molecular detection.

The manuscript was written according to the STROBE statement guidelines.

## RESULTS

### Study Population

A total of 600 pregnant women were enrolled in a prospective cohort study (PCS) and followed until delivery. Of the first eligible maternal-child pairs, 409 (68.2%) met the inclusion criteria (Fig. 2). Among the 409 newborns, 251 were born from mothers that had malaria infection during pregnancy, *P. vivax* (*Pv*), *P. falciparum* (*Pf*) or both (mixed) (Fig. 2). Overall, there were no relevant maternal and newborns baseline differences between the distinct groups (Additional file 2). Nonetheless, women that were *Plasmodium*-infected presented few characteristics at delivery that were slightly different from the Non-Infected group: less weight gain, lower hematocrit, lower hemoglobin, and reduced placental weight (Additional file 2).

**Figure 2.**
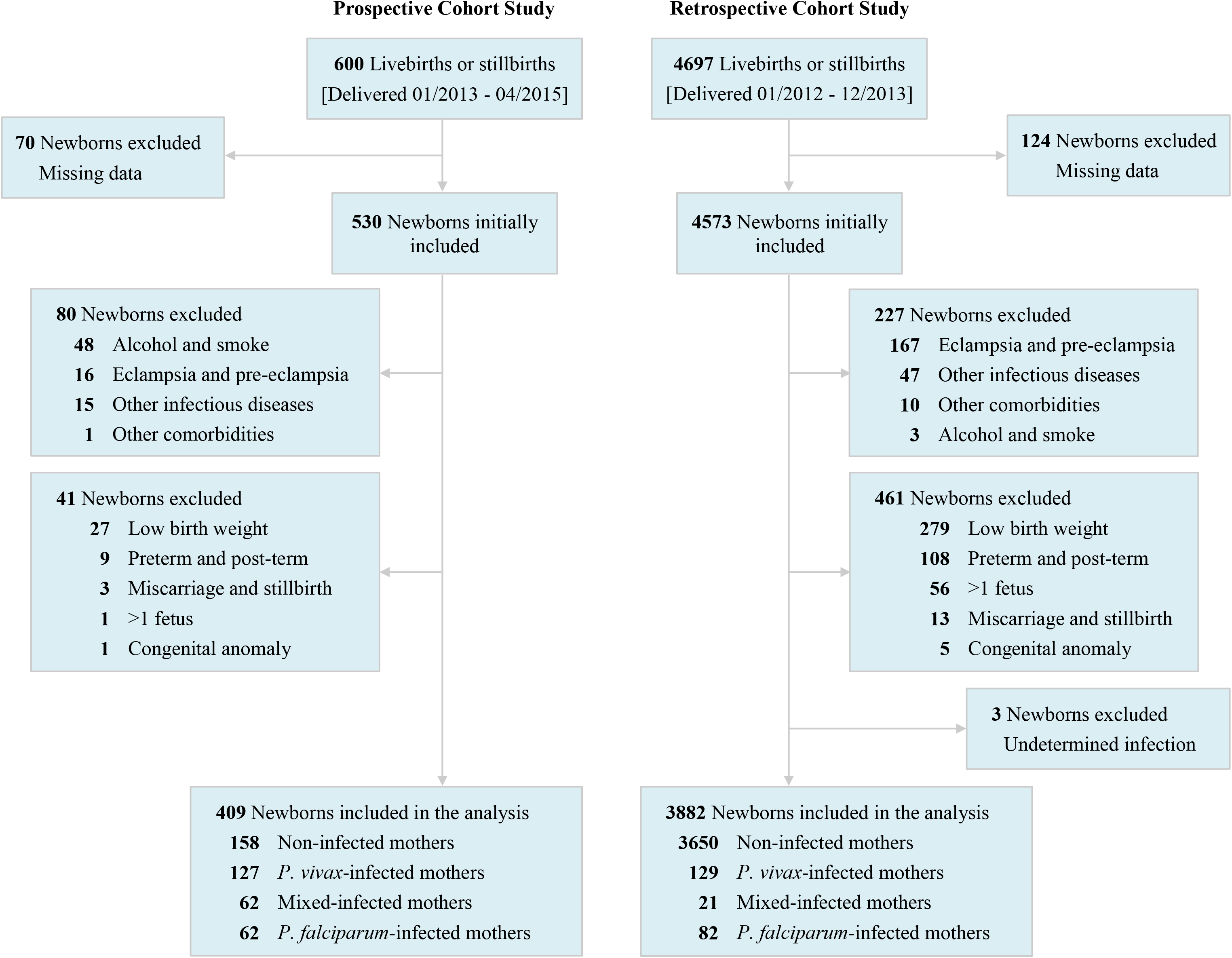
Flow diagram of the two cohort studies detailing exclusion criteria. Mixed infection – *P. vivax*- and *P. falciparum*-infection occurring at the same time and/or at different times during pregnancy.

### Reduced head circumference in newborns from women infected with *P. falciparum* during pregnancy

The frequency distribution of the newborns HC born from non- (NI) and malaria-infected mothers (Malaria), including LBW and preterm babies, evidenced differences between the two groups. The Malaria group displayed a deviated peak and spread to the left when compared with the NI group, indicative of more newborns with reduced HC (*p* = 0.005) (Fig. 3a). Nevertheless, to assure that the observed difference was not due to the LBW and preterm babies, these newborns were removed from the analysis and segregated the malaria-infected group into *Plasmodium* species infected groups. Even though, it was possible to observe an apparent deviation of the peak of the *P. falciparum-infected* group (*Pf*) from the non-infected (NI) (*p* = 0.023) (Fig. 3b), indicating a higher frequency of babies with smaller HC when mothers are infected by *P. falciparum*.

**Figure 3.**
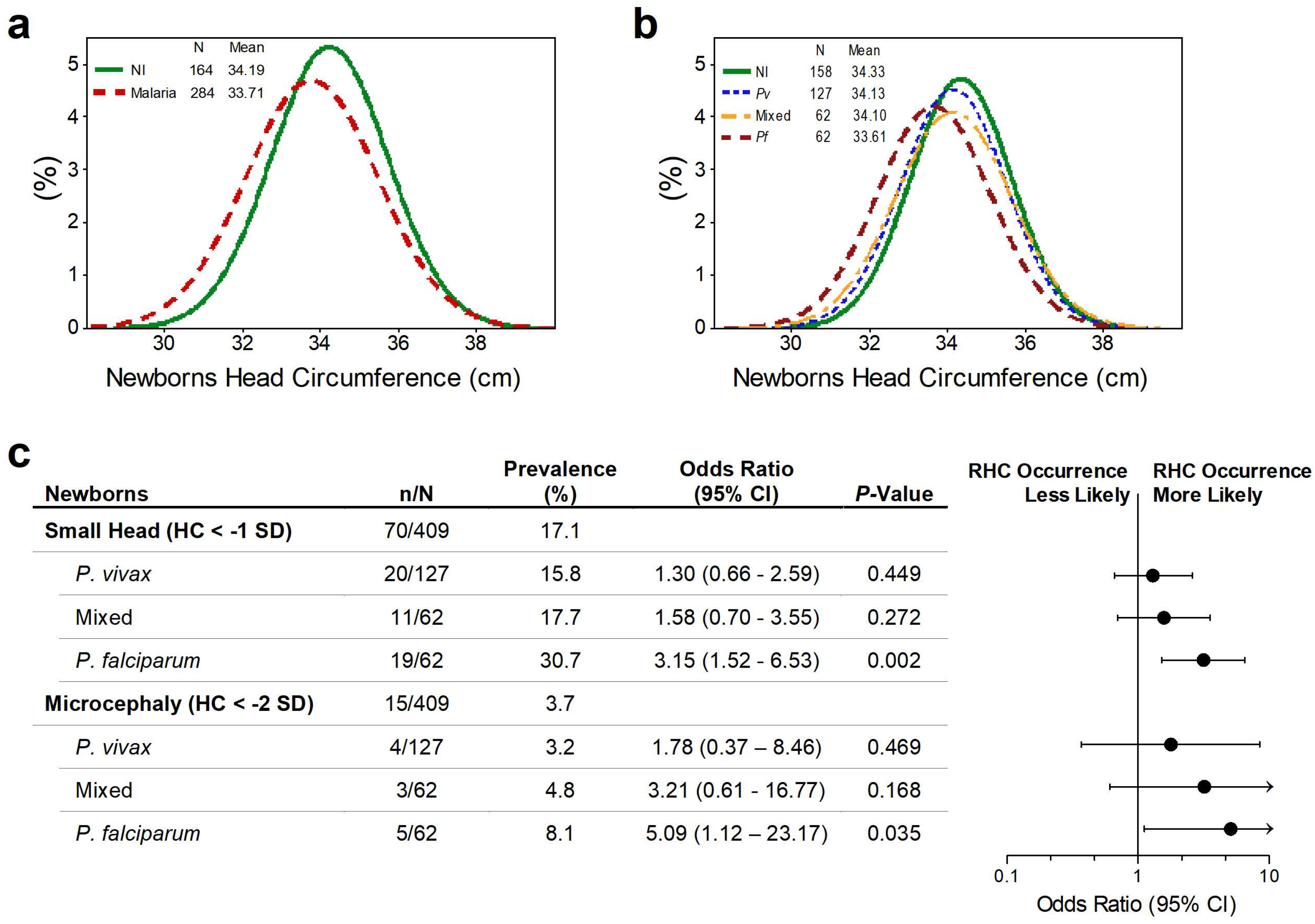
Prospective cohort study shows that malaria infection during pregnancy impacts babies head circumference. **a, b** Newborns head circumference frequency distribution in the PCS according to maternal infection status: malaria- and non-infected (NI) mothers (*p* = 0.005) (a), and NI, *Pv*, Mixed and Pf-infected mothers after excluding LBW and preterm babies (NI vs *Pf p* = 0.023) (**b**). The differences in the frequency distributions between each group were examined with Mann-Whitney rank sum tests. c Forest plot of the Odds Ratio of small head or microcephaly in babies born from women infected during pregnancy compared to babies from non-infected women, according to *Plasmodium* species. Mixed infection – *P. vivax-* and *P. falciparum*-infection occurring at the same time and/or at different times during pregnancy. n/N - number of events by total number of individuals in each group; CI - confidence interval; HC - head circumference; SD - standard deviation; *P*-Values were estimated through multivariate logistic regression methods.

Among the evaluated newborns in the PCS, 70 (17.1%) babies presented with a small head (SH), including 15 (3.7%) with microcephaly (Fig. 3c). The evaluated babies were considered proportionate through the Rohrer Index, independently of the HC size (Additional file 3). Further, to evaluate the association of malaria during pregnancy with fetus head growth, the newborns were segregated by HC and the mother infection status: non-infected, *P. vivax-*, mixed- or *P. falciparum*-infected. The prevalence of newborns with SH was higher among babies born from women infected with *P. falciparum* (30.7%) during pregnancy. Similarly, the prevalence of microcephaly doubled when a *P. falciparum* infection has occurred (8.1%) (Fig. 3c). In fact, a multivariate logistic regression analysis identified *P. falciparum* infection as increasing the odds of occurring SH in newborns (OR 3.15, 95% CI 1.52-6.53, *p* = 0.002) (Fig. 3c). Likewise, it revealed a higher likelihood of occurring microcephaly in babies born from mothers that were *P. falciparum-infected* (OR 5.09, 95% CI 1.12-23.17, *p* = 0.035) (Fig. 3c). Strikingly, *P. vivax* infection during pregnancy was not found to be associated with reduced HC (for SH, OR 1.30, 95% CI 0.66-2.59, *p* = 0.449). Maternal-child pairs that presented misleading factors such as TORCH infections, Syphilis, HIV, Dengue, Chikungunya, and Zika virus, and alcoholism and drug use declared in the medical records, or identified in all mothers that delivered babies with were discarded SH (Additional file 4).

### Reduced head circumference in newborns is associated with placental malaria

Further, several placental parameters were evaluated to ascertain the relation of placental malaria due to *P. falciparum* infection with the SH occurrence. Strikingly, babies with SH (Pf-SH) born from mothers that had their first infection later in gestation (median [IQR], 25.5 weeks [18.0-32.5], *p* = 0.014) when compared with NHC (19.0 weeks [12.0-29.3]). Moreover, much of the placental malaria manifestation in newborns with SH (Pf-SH) or microcephaly (Pf-MC) was due to a past *P. falciparum* infection (54% and 72%, respectively), as opposed to 48% in placentas from newborns with NHC (Pf-NHC) (Table 1).

**Table 1.** Infection characteristics in *P. falciparum-infected* pregnant women.

|  | *Pf*-NHC<br>(N=94) | *Pf*-SH<br>(N=30) | *p*-Value<sup>a</sup> | *Pf*-MC<br>(N=8) | *p*-Value<sup>b</sup> |
| --- | --- | --- | --- | --- | --- |
| Infections per pregnancy, median (IQR) | 2.0 (1.0-3.0) | 2.0 (1.0-2.0) | 0.463 | 1.0 (1.0-2.0) | 0.116 |
| Parasitemia of first infection, median (IQR)<sup>c</sup> | 1.2 (0.3-4.6) | 3.8 (0.5-9.2) | 0.053 | 0.4 (0.2-1.8) | 0.220 |
| Gestational age at first infection |  |  |  |  |  |
| Mean (SD) | 20.7 (10.5) | 26.0 (8.1) | 0.014 | 27.6 (7.8) | 0.064 |
| Median (IQR) | 19.0 (12.0-29.3) | 25.5 (18.0-32.5) |  | 28.5 (19.8-34.3) |  |
| Placental Malaria, no. (%)<sup>d</sup> |  |  |  |  |  |
| No | 29 (36) | 7 (30) | - | 1 (14) | - |
| Active Acute | 8 (10) | 2 (8) | - | 0 | - |
| Active Chronic | 5 (6) | 2 (8) | - | 1 (14) | - |
| Past | 38 (48) | 13 (54) | - | 5 (72) | - |
| Hemozoin, no. (%)<sup>d, e</sup> |  |  |  |  |  |
| No | 31 (39) | 8 (33) | - | 1 (14) | - |
| Mild | 32 (40) | 9 (38) | - | 4 (57) | - |
| Moderate | 15 (19) | 7 (29) | - | 2 (29) | - |
| Severe | 2 (2) | 0 | - | 0 | - |

N, number of individuals; *Pf*-NHC, *Plasmodium falciparum*-normal head circumference; *Pf*-SH,
*Plasmodium falciparum*-small head; *Pf*-MC, *Plasmodium falciparum*-microcephaly; IQR,
interquartile range; SD, standard deviation; no., number of events.
<sup>a</sup> Differences between *Pf*-NHC and *Pf*-SH groups were evaluated using Mann-Whitney rank sum tests.
<sup>b</sup> Differences between *Pf*-NHC and *Pf*-MC groups were evaluated using Mann-Whitney rank sum tests.
<sup>c</sup> Parasitemia was recorded in 82 *Pf*-NHC, 28 *Pf*-SH, and 7 *Pf*-MC. Values presented in 10<sup>3</sup> DNA copies, obtained by PET-PCR quantification.
<sup>d</sup> Placental malaria and Hemozoin was recorded in 80 *Pf*-NHC, 24 *Pf*-SH, and 7 *Pf*-MC.
<sup>e</sup> Hemozoin - Mild: focal presence in small amounts; Moderate: small spots or larger deposits in many locations; Severe: large amounts present widely.

The analysis of placental histology parameters and angiogenic factors disclosed substantial differences between non-infected controls and *P. falciparum-infected* groups. Of note, in all *P. falciparum-infected* groups, we observed higher monocytes infiltrate (median[IQR], Pf-NHC 7.0 [5.0-13.0], *p* <0.0001; Pf-SH 9.5 [5.5-15.0], *p* <0.0001; Pf-MC 9.0 [6.0-11.0], *p* = 0.018 vs Non-Infected 4.0 [2.0-7.0]) (Fig. 4c, d). On the other hand, the syncytial nuclear aggregates (SNA) and Leptin alterations were only observed in infected placentas of babies with SH and MC. Remarkably, SNA that have a long-standing association with placental pathologies (Heazell et al. 2007), presented excessive formation in the Pf-SH and Pf-MC groups (17.5 [12.0-24.5], *p* = 0.002 and 18.0 [12.0-30.0], *p* = 0.023, respectively) when compared to the Non-Infected (13.0 [10.0-17.0]) (Fig. 4g, h), as well, when Pf-SH was compared to Pf-NHC. Moreover, the Leptin levels were markedly reduced in the Pf-SH and *Pf-MC* groups (19.5 [4.5-37.2], *p* = 0.013 and 16.7 [9.0-26.7], *p* = 0.027, respectively) when compared to the Non-Infected (33.1 [17.2-47.4]) (Fig. 5i). Complete data details can be found in Additional file 5.

**Figure 4.**
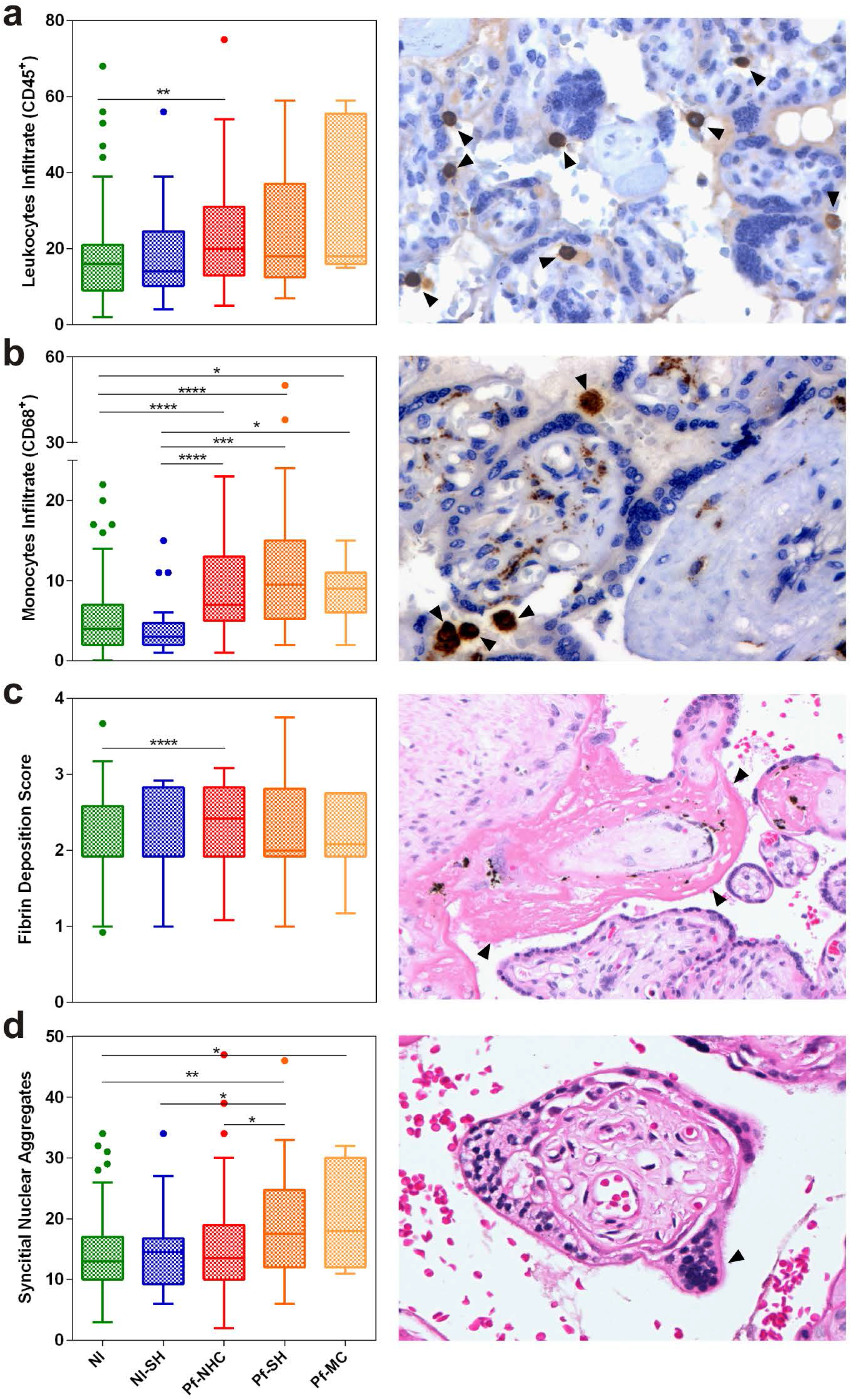
Histopathological parameters evaluation of placentas from non- and *P. falciparum*-infected mothers according to newborns head circumference. **a** Leukocytes (CD45^+^) number. **b** Monocytes (CD68^+^) number. **c** Fibrin deposition score. **d** Syncytial nuclear aggregates. Images in each panel are only representative. Histopathological parameters were evaluated by microscopy through H&E (fibrin deposition and syncytial nuclear aggregates) and immunohistochemistry (leukocytes and monocytes) staining. NI - non-infected; NI-SH - non-infected small head; Pf-NHC - *P. falciparum-infected* normal head circumference; Pf-SH - *P. falciparum*-infected small; and, Pf-MC - *P. falci*parum-infected microcephaly. Data are represented as Tukey boxplots, the bottom and the top of the box are the first and third quartiles, the line inside the box is the median, and the whiskers represent the lowest and the highest data within 1.5 IQR of the first and upper quartiles. The differences between each group were examined with Mann-Whitney rank sum tests, * *p* < 0.05, ** *p* < 0.01, *** *p* < 0.001 and **** *p* < 0.0001.

**Figure 5.**
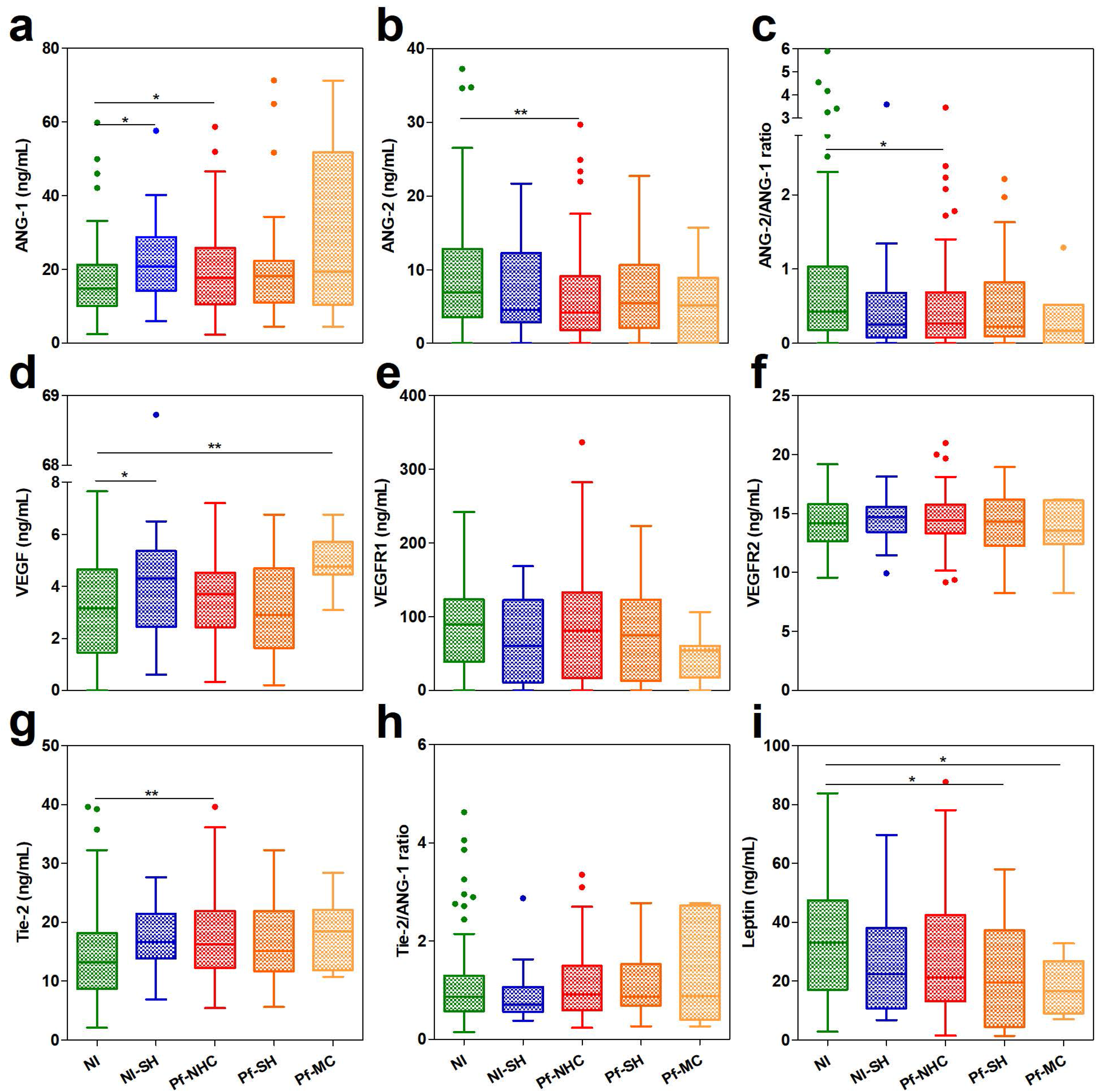
Placental plasma levels of angiogenic factors and leptin from non- and *P. falciparum-infected* mothers according to newborns head circumference. **a** Angiopoietin-1 (ANG-1). **b** Angiopoietin-2 (ANG-2). **c** Ratio ANG-2/ANG-1. **d** Vascular endothelial growth factor (VEGF). **e** VEGF receptor-1 (VEGFR-1). **f** VEGF receptor-2 (VEGFR-2). **g** TEK receptor tyrosine kinase (Tie-2). **h** Ratio Tie-2/ANG-2. i Leptin. All factors were measured by ELISA. NI – noninfected; NI-SH – non-infected small head; Pf-NHC – *P. falciparum*-infected normal head circumference; Pf-SH – *P. falciparum*-infected small; and, Pf-MC - *P. falci*parum-infected microcephaly. Data are represented as Tukey boxplots, the bottom and the top of the box are the first and third quartiles, the line inside the box is the median, and the whiskers represent the lowest and the highest data within 1.5 IQR of the first and upper quartiles. The differences between each group were examined with Mann-Whitney rank sum tests, * *p* ≤ 0.05, ** *p* < 0.01.

Furthermore, evaluation of inflammatory factors in the placental plasma revealed differences mainly between the Non-Infected group and the Pf-NHC group. Though, the Pf-SH group shows statistically significant higher IL8 and smaller C3a plasma levels (45.1 [22.1-85.9], *p* = 0.044; and, 3.0 [0-5.5], *p* = 0.014, respectively) when compared to the Non-Infected group (25.5 [15.7-52.2], and, 4.5 [3.2-6.6], respectively) (Additional file 5). These results support a placental dysfunction upon *P. falciparum* infection, which in some parameters are specifically heightened in placentas derived from babies with reduced HC, like the syncytial nuclear aggregates.

### Retrospective cohort study corroborates the reduced head circumference association with *P. falciparum* infection

Further, a population-based retrospective cohort study (RCS) was conducted to confirm the association results. A total of 4697 maternal-child pairs were included, and upon application of the exclusion criteria, 3882 (83%) newborns remained to be evaluated, of which, 232 were born from mothers that had malaria infection during pregnancy (Fig. 2). Overall, there were no significant differences in baseline characteristics between the PCS and the RCS (Additional file 2 and 6). The evaluation of the frequency distribution of the newborns HC born from non-(NI) and malaria-infected mothers (Malaria), showed differences between the two groups *(p* = 0.008) (Fig. 6a). Identical to the PCS, when the LBW and preterm babies were removed from the analysis, and the malaria-infected group segregated, the *P. falciparum-infected* group *(Pf)* presented a deviated peak from the non-infected (NI) *(p* = 0.015) (Fig. 6b). Indicative of a higher frequency of newborns with reduced HC when mothers are infected with *P. falciparum* during pregnancy.

**Figure 6.**
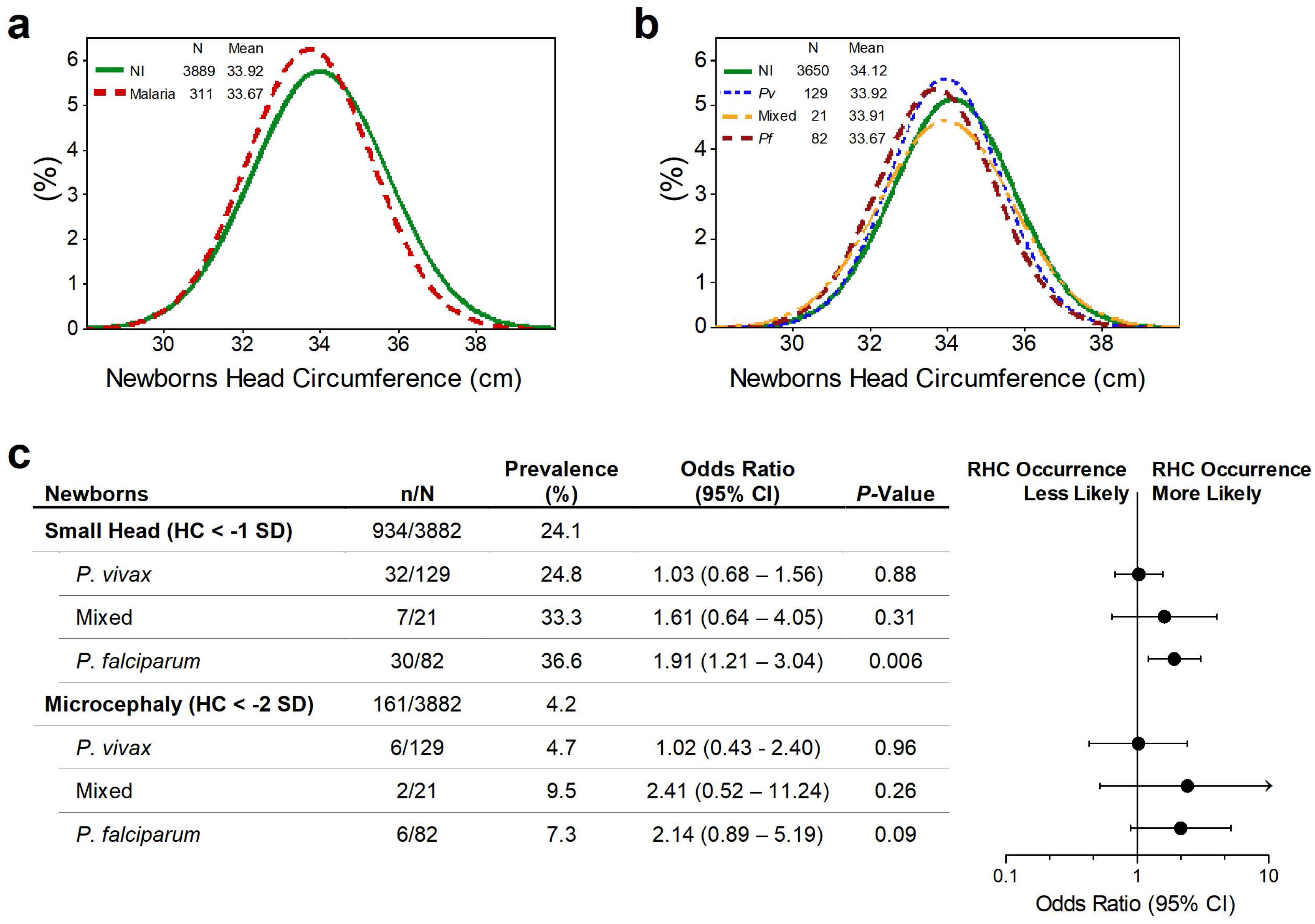
Retrospective cohort study corroborates that malaria infection during pregnancy impacts babies head circumference. **a**, **b** Newborns head circumference frequency distribution in the PCS according to maternal infection status: malaria- and non-infected (NI) mothers (*p* = 0.008) (**a**), and NI, *Pv*, Mixed and Pf-infected mothers after excluding LBW and preterm babies (NI vs *Pf p* = 0.015) (**b**). The differences in the frequency distributions between each group were examined with Mann-Whitney rank sum tests. c Forest plot of the Odds Ratio of small head or microcephaly in babies born from women infected during pregnancy compared to babies from non-infected women, according to *Plasmodium* species. Mixed infection – *P. vivax*- and *P. falciparum*-infection occurring at the same time and/or at different times during pregnancy. n/N - number of events by total number of individuals in each group; CI - confidence interval; HC - head circumference; SD - standard deviation; *p*-Values were estimated through multivariate logistic regression methods.

The evaluated newborns included 934 (24.1%) babies with SH and 161 (4.2%) with microcephaly. In the RCS, similarly to the PCS, the prevalence of newborns with SH was more than one-half higher (36.6%) among babies born from *P. falciparum-infected* mothers, and the microcephaly prevalence almost doubled in the presence of a *P. falciparum* infection (7.3%) (Fig. 6c). Analogously, the multivariate logistic regression analysis revealed that *P. falciparum* infection increases the odds of occurring SH in newborns (Odds ratio [OR] 1.91, 95% CI 1.21-3.04, *p* = 0.006) (Fig. 6c). Altogether, these results demonstrate that *P. falciparum* infection during pregnancy increases the likelihood of occurring reduced HC in the newborns, corroborating the results obtained in the PCS.

## DISCUSSION

It is well-established that malaria during pregnancy increases the risk of adverse fetal outcomes, such as abortion, IUGR, premature births and LBW. We show evidence that *P. falciparum* infection during pregnancy is significantly associated with the occurrence of reduced HC in the newborns, and to some extent, with microcephaly. The revealed newborn HC reduction is independent of the already known impact that malaria has on the whole fetal growth, as LBW and preterm newborns were deliberately excluded from our analysis.

The increased risk for developing reduced HC associated with *P. falciparum* infection was supported by a prospective study (PCS) (Odds Ratio (OR) 3.15, *p* = 0.002) and subsequently corroborated by a retrospective study (RCS) (OR 1.91, *p* = 0.006). Remarkably, in the prospective study, the OR doubles when we consider only the microcephaly cases (OR 5.09, *p* = 0.035). These observations reinforce the knowledge that malaria during pregnancy increases the risk of problems in fetal development (Desai et al. 2007; Ismail et al. 2000; Rogerson et al. 2007).

We hypothesize that the placental inflammatory process acting upon *P. falciparum* infection is contributing to impair the fetal head growth. This hypothesis is supported by the observation of histopathological alterations, combined with an imbalance in angiogenic factors production and inflammatory factors in placentas from babies with congenital SH or microcephaly when mothers were *P. falciparum-infected*. A local inflammation can generate a frame of hypoxia/ischemia that would alter the transportation of both nutrients and respiratory gases to the unborn baby, which can impact on cranial malformation due to the lack of an adequate supply of nutrients and oxygen (Nelson and Penn 2015). Also, the oxidative stress caused by hypoxia leads to several structural and functional alterations in the intrauterine development (Kurinczuk, White-Koning, and Badawi 2010). This scenario is often observed in cases of placental malfunction due to different etiologies, and prolonged and premature labor (Boksa 2004).

Interestingly, the values of SNA or syncytial knotting, which has been associated with IUGR due to local hypoxia/oxidative stress (Heazell et al. 2007), were highly increased in placentas from the *Pf*-SH and *Pf*-MC groups when compared to the other control groups. Syncytial knotting has repeatedly been observed in placentas from *P. falciparum-exposed* women (Souza et al. 2013; Bulmer et al. 1993; Ismail et al. 2000). In fact, the major placental alterations observed, including syncytial knots and monocytes inflammatory infiltrate, are consistent with previous reports on placental inflammatory responses due to sequestration of *P. falciparum* parasites in the placenta, which characterizes the placental malaria development (Ismail et al. 2000; Rogerson et al. 2007; Souza et al. 2013). The evaluation of cytokine levels and complement in our samples did not show an overall alteration. Nevertheless, these only reflect a picture at the moment of birth. It is unsurprising that *P. vivax* infection was not associated with the head reduction phenotype, as this parasite is known as not sequestering in the placenta. Previous studies have demonstrated that *P. vivax* infection during pregnancy induces a less placental inflammatory process when compared with *P. falciparum* infection (Souza et al. 2013).

The presence of residual tissue lesions and impaired leptin production constitute clear evidence of damage. In fact, the Pf-SH and Pf-MC groups presented deregulated leptin levels. The impaired production of leptin, a hormone commonly produced in substantial amounts by the placenta, can be related to placental inflammation upon infection. Also, leptin has been shown associated with fetal growth restriction (Conroy et al. 2011). Regarding the Pf-SH group, few observed differences reached statistical significance, possibly due to the small sample size of this group, but the overall placental malaria phenotype is more prominent and widespread than in non-infected and Pf-NHC groups. Nevertheless, it is unclear how placental alterations due to inflammation impact on the development of the fetus.

Currently, much of what is known about falciparum gestational malaria is based on studies performed in African high transmission areas, which in general are settings that have precarious health systems and inadequate or late treatment provision. In Brazil, approximately 85% of the infections are caused by *P. vivax. P. falciparum* is only transmitted in specific regions, including in the one evaluated in this work (“Alto do Juruá” valley, Acre), where it is responsible for 46% of the total infections in Brazil (SIVEP - Secretaria de Vigilância em Saúde - Ministério da Saúde 2015; Ferreira and Castro 2016). Interestingly, despite Brazil being a low transmission area for malaria with effective control strategies and early treatment provision, we observed adverse events in newborns similar to those reported in areas of high endemicity.

Surprisingly, the prevalence of microcephaly (HC < -2 SD) observed by us is far higher than what has been previously reported by the Brazilian Ministry of Health (Passemard, Kaindl, and Verloes 2013). Two independent studies have recently evaluated retrospectively babies born in two different Brazilian regions, and also reported a higher prevalence of microcephaly in babies born before the Zika outbreak (Soares de Araújo et al. 2016; Magalhães-Barbosa et al. 2017). In one, 16,208 infants born between 2012 and 2015 in the Paraiba State (Brazil) were evaluated, and 4.2 to 8.2% of microcephaly prevalence was reported, depending on the classification criteria (Soares de Araújo et al. 2016). In the other, 8,275 babies born between 2011 and 2015 in the southeastern and mid-western Brazilian region were evaluated, and an overall prevalence of microcephaly of 5.6% was identified (Magalhães-Barbosa et al. 2017). In fact, it is puzzling that a country like the USA with about 3.5 millions of births per year reports annually approximately 25,000 infants with microcephaly (Ashwal et al. 2009); on the other hand, Brazil with about 3 million births per year reported around 150 microcephaly cases annually, before Zika epidemy (Ministério da Saúde - Secretaria de Vigilância em Saúde-Brasil 2015). These observations indicate an inconsistency of the data released by the Brazilian authorities probably due to under-reporting.

Our work has some potential limitations. First, the babies’ HC was only assessed at birth, since it was not possible to perform the morphometric measures through ultrasonography during pregnancy in the public health system, as well as the possibility of acquiring newborn head imaging. Second, reduction of HC has different etiologies, namely, genetic causes and action of infectious agents. While we have discarded misleading factors, such as TORCH infections, Syphilis, HIV, Dengue, Chikungunya and Zika virus, as well as smoking, alcoholism and drug use, studies to detect genetic abnormalities in those patients were not performed. Third, although in both the PCS and the RCS the logistic-regression analysis indicates a strong association between SH and *P. falciparum* infection, we only had access to few placentas. The smaller sample size has limited the statistical analysis; however, most of the parameters analyzed indicated intensified placental malaria when compared to placentas from newborns with normal head size.

## CONCLUSION

This work provides evidence that *P. falciparum* infection during pregnancy can impact the head growth of the fetus, which leads to small heads and in extreme cases to microcephaly. If our results are confirmed, the consequences of gestational malaria over fetal neurological development, which can lead to poor neurocognitive and behavioral development, represents a serious long-term health problem. Physicians should periodically assess the development and academic achievements of these children, with a comprehensive neurocognitive evaluation, to guide preventive and rehabilitative assistance that might improve outcomes. Extensive epidemiological prospective studies, involving the collection of biological, clinical, and socioeconomic data and potential confounding factors, are required to establish the prevalence of SH and microcephaly and its association with malaria. Our work reinforces the urgent need to protect the pregnant women and their unborn babies from the devastating effects of malaria infection.

## ABBREVIATIONS

ANC: : Antenatal care;
ANG-1 and ANG-2: : Angiopoietins 1 and 2;
CI: : Confidence intervals;
cm: : centimeters;
g: : Grams;
HC: : head circumference;
H&E: : Hematoxylin-Eosin;
HMCJ: : Hospital da Mulher e da Criança do Juruá;
IUGR: : Intrauterine growth retardation;
IQR: : Interquartile ranges;
LBW: : Low birth weight;
LMP: : Last menstrual period;
MC: : Microcephaly;
MoH: : Ministry of Health;
NHC: : Normal head circumference;
NI: : Non-infected;
OD: : Optical density;
OR: : Odds ratio;
PCS: : Prospective cohort study;
Pf: : *Plasmodium falciparum;*
Pv: : *Plasmodium vivax;*
RCS: : Retrospective cohort study;
SD: : standard deviations;
SH: : Small head;
SIVEP: : Epidemiological Surveillance Information System;
SNA: : Syncytial nuclear aggregates;
TIE-2: : TEK receptor tyrosine kinase;
TMA: : Tissue microarray;
TORCH: : abbreviation for Toxoplasma, rubella, cytomegalovirus, and Herpes simplex;
VEGFA: : Vascular endothelial growth factor A;
WHO: : World Health Organization;
WHO-CGS: : WHO child growth standards.

## DECLARATIONS

### Ethics approval and consent to participate

Ethical clearance was provided by the committees for research of the University of São Paulo and the Federal University of Acre (Plataforma Brasil, CAAE: 03930812.8.0000.5467 and 03930812.8.3001.5010, respectively), according to Resolution n° 196/96 of Brazilian National Health Committee. All the study participants or their legal guardians (if minors) gave written informed consent. The authors have agreed to maintain the confidentiality of the data collected from the medical records and databases, by signing the Term of Commitment for the Use of Data from Medical Records. The study was conducted in accordance with the Declaration of Helsinki and is registered in the Brazilian Clinical Trials Registry as RBR-3yrqfq.

### Consent for publication

Not applicable.

### Availability of data and materials

All relevant data are available from the authors on request.

### Competing interests

The authors declare that they have no competing interests.

### Funding

This work was primarily funded by grants from São Paulo Research Foundation (FAPESP), CRFM (2009/53889-0 and 2014/09964-5) and SE (2014/20451-0). JMS was supported by CNPq (308613/2011-2) and FAPESP (2013/21728-2). PMAZ was supported by FAPESP (2014/17766-9). MAGG was supported by CNPq (404478/2012-3). TGC was supported by the Medical Research Council UK (Grant no. MR/K000551/1, MR/M01360X/1, MR/N010469/1, MC_PC_15103). SC was funded by the Medical Research Council UK (Grant no. MR/M01360X/1, MC_PC_15103). JGD, FAL, OM, MPC, and LAG were supported by FAPESP fellowships (2012/04755-3, 2013/16417-8, 2013/00981-1, 2016/08204-2, and 2015/06106-0, respectively). The funders had no role in analysis design, data collection and analysis, decision to publish, or preparation of the manuscript.

### Authors’ contributions

JGD, RMS, SE, and CRFM designed the study. JGD, RMS, FAL, CLB, OM, DSC, EPMP, MPC, PMAZ, MAGG, SE, LAG, and CRFM were involved in data acquisition and scientific input. JGD, RMS, FAL, CLB, OM, DSC, EPMP, MPC, PMAZ, EB, MAGG, SC, TGC, SE, LAG, and CRFM contributed to the analysis and/or interpretation of data. ACPL, JMS, and TGC performed the multivariate logistic regression analysis. LAG and CRFM wrote the manuscript and compiled the information in the Additional information. CRFM and SE were the main funders of this work. CRFM have had full access to all the data in the study and takes responsibility for the integrity of the data and the accuracy of the data analysis. All authors reviewed and approved the final version of this manuscript.

## Acknowledgements

We thank the women from “Alto do Juruá” valley who agreed to participate in the study, as well as the nurses and technicians from the Hospital da Mulher e da Criança do Juruá and Gerência de Endemias/SESACRE team for their invaluable assistance. Also, we thank the direction of Santa Casa de Misericórdia de Cruzeiro do Sul, and Universidade Federal do Acre for the support. Additionally, we thank Alexandre Macedo de Oliveira from Centers for Disease Control and Prevention (CDC) for his ongoing support of our study; Ricardo Ataíde for assistance during fieldwork and scientific input; and Bernardo Paulo Albe for technical assistance. Finally, we thank Venkatachalam Udhayakumar, Luciana Flannery and Naomi Lucchi from Malaria Laboratory Research and Development Unit at CDC for all the support on the establishment and training of the PET-PCR technique, which was funded by the U.S. Agency for International Development (USAID) through the Amazon Malaria Initiative (AMI).

## ADDITIONAL FILE

**Additional file 1:** Summary of histopathological evaluation methods. (PDF)

**Additional file 2:** Summary of maternal and newborns characteristics of the Prospective Cohort Study (PCS). (PDF)

**Additional file 3:** Summary of newborns characteristics of the Prospective Cohort Study according with head circumference. (PDF)

**Additional file 4:** Summary of serological screening of TORCH, HIV, Syphilis, Dengue, Chikungunya and Zika infections. (PDF)

**Additional file 5:** Summary of placental parameters evaluation in the Prospective Cohort study according to newborns head circumference. (PDF)

**Additional file 6:** Summary of maternal and newborns characteristics of the Retrospective Cohort Study (RCS). (PDF)

---

**Additional file 1: Summary of histopathological evaluation methods.**

**Table 1.** Evaluation Methods and Staining Used to Quantify Malaria-Associated Placental Parameters

|  | Evaluation methods | Staining |
| --- | --- | --- |
| **Pathological features** |  |  |
| Syncytial nuclear aggregates | Number of affected villi per 100 villi at 10× magnification.[1,2] | Hematoxylin-Eosin |
| Fibrin deposition score | Semi-quantitative scoring was used for placental fibrin on a scale from 0 to 5. For the extent of fibrin deposition at the basal and chorionic plates and intervillous fibrin and perivillous fibrin, the following scale was used to apply a score to each: none (0), scant (1), minimal extension (2), moderate (3), heavy (4), or extensive (5) at 100× magnification.[1,2] | Hematoxylin-Eosin |
| Fibrinoid Necrosis | Number of intersection points on a random grid that touched areas of necrosis per total points of a square grid 4 862.43 $\mu\text{m}^2$ of area point at 10× magnification.[1,2] | Hematoxylin-Eosin |
| Proliferation index | The index was calculated by quantitative image analysis of the percentage of positively stained nuclear area with anti-Ki-67/DAB per the total area of the nuclei, obtained by averaging three images of the same sample at 20-fold increase. Employing a web available free application for ImageJ.[3] | Immunohistochemistry |
| Villous vascularity | Number of fetal vessels labeled with anti-CD31 in ten villi terminals at 20× magnification in the Axio Scan.Z1 scanning system. | Immunohistochemistry |
| Leukocytes infiltrate (CD45<sup>+</sup>) | Number of leukocyte cells in 10 fields at 400× magnification. | Immunohistochemistry |
| Monocytes infiltrate (CD68<sup>+</sup>) | Number of monocyte cells in 10 fields at 400× magnification.[4–6] | Immunohistochemistry |
| **Malaria-associated features** |  |  |
| Parasitized erythrocytes | Number of fields with parasite in 100 fields at 1000× magnification. | Giemsa |
| Hemozoin | Sixty fields at 40× magnification were screened with polarized light for the presence of hemozoin in the intervillous space (free or within cells) and in the tissue.[7] | Hematoxylin-Eosin |

**Additional file 2: Summary of maternal and newborns characteristics of the Prospective Cohort Study (PCS).**

**Table 1.** Baseline characteristics of mothers

| Mothers' Characteristics | Non-Infected<br>(N=158) | *P. vivax*<br>(N=127) | Mixed<br>(N=62) | *P. falciparum*<br>(N=62) |
| --- | --- | --- | --- | --- |
| Maternal age (years), mean (SD) | 24.3 (6.2) | 22.2 (6.2) | 23.3 (5.8) | 23.1 (6.3) |
| Gravidity, no. (%) |  |  |  |  |
| Primigravida | 72 (45.6) | 51 (40.2) | 19 (30.7) | 23 (37.1) |
| Multigravida | 86 (54.4) | 76 (59.8) | 43 (69.3) | 39 (62.9) |
| Gestational age at delivery (weeks) |  |  |  |  |
| Mean (SD) | 39.7 (1.2) | 39.6 (1.3) | 39.5 (1.2) | 39.6 (1.3) |
| Median (IQR) | 40.0 (39.0-40.0) | 40.0 (39.0-41.0) | 40.0 (39.0-40.0) | 39.5 (39.0-40.0) |
| C-section, no. (%) | 90 (57.0) | 51 (40.2) | 27 (43.6) | 23 (37.1) |
| Maternal weight gain (Kg), mean (SD)<sup>a</sup> | 13.6 (5.0) | 11.2 (5.0) | 12.0 (5.0) | 11.8 (5.2) |
| Hematocrit (%), mean (SD)<sup>b</sup> | 36.2 (3.5) | 35.3 (3.9) | 34.9 (4.0) | 34.2 (4.3) |
| Hemoglobin (g/dL), mean (SD)<sup>c</sup> | 11.9 (1.2) | 11.6 (1.3) | 11.5 (1.3) | 11.2 (1.4) |
| Placental weight (g), mean (SD)<sup>d</sup> |  |  |  |  |
| Primigravida | 578.8 (97.0) | 558.7 (102.2) | 533.3 (63.1) | 568.0 (129.2) |
| Multigravida | 608.0 (112.2) | 601.5 (148.9) | 578.8 (108.8) | 592.6 (159.1) |
| Antenatal care visits, mean (SD)<sup>e</sup> | 7.9 (2.3) | 6.4 (2.5) | 6.3 (2.3) | 5.6 (2.8) |
| Previous malaria episodes during current pregnancy, no. (%) | - | 37 (29.1) | 30 (48.4) | 7 (11.3) |

N, number of individuals; SD, standard deviation; IQR, interquartile range; no., number of events.
<sup>a</sup> Maternal weight gain was recorded in 153 non-infected, 107 *P. vivax*, 56 mixed-infected and 49 *P. falciparum* pregnant women. It was determined by subtracting the initial pregnancy weight from the final weight.
<sup>b</sup> Hematocrit was recorded in 107 non-infected, 86 *P. vivax*, 43 mixed-infected and 35 *P. falciparum* pregnant women.
<sup>c</sup> Hemoglobin was recorded in 107 non-infected, 85 *P. vivax*, 43 mixed-infected and 35 *P. falciparum* pregnant women.
<sup>d</sup> Placental weight was recorded in 148 non-infected, 108 *P. vivax*, 57 mixed-infected and 48 *P. falciparum* pregnant women.
<sup>e</sup> The number of antenatal care visits was recorded in 153 non-infected, 120 *P. vivax*, 59 mixed-infected and 57 *P. falciparum* pregnant women.

**Table 2.** Baseline characteristics of newborns

| Newborns' Characteristics | Median (IQR) |  |  |  |
| --- | --- | --- | --- | --- |
|  | Non-Infected<br>(N=158) | *P. vivax*<br>(N=127) | Mixed<br>(N=62) | *P. falciparum*<br>(N=62) |
| Male newborns, no. (%) | 72 (45.6) | 68 (53.5) | 35 (56.5) | 29 (46.8) |
| Weight (g) |  |  |  |  |
| Male | 3250.0 (3012.5-3477.5) | 3360.0 (3067.5-3625.0) | 3250.0 (3015.0-3420.0) | 3320.0 (3055.0-3540.0) |
| Female | 3365.0 (3045.0-3630.0) | 3065.0 (2865.0-3460.0) | 3060.0 (2890.0-3400.0) | 3115.0 (2935.0-3260.0) |
| Length (cm)<sup>a</sup> |  |  |  |  |
| Male | 49.0 (48.0-50.0) | 49.0 (49.0-50.0) | 50.0 (48.0-50.0) | 49.0 (48.0-50.0) |
| Female | 49.5 (49.0-50.0) | 49.0 (48.0-50.0) | 49.0 (48.0-50.0) | 49.0 (48.0-50.0) |
| Rohrer index<sup>a,b</sup> |  |  |  |  |
| Male | 2.7 (2.5-2.9) | 2.7 (2.6-2.9) | 2.7 (2.5-2.9) | 2.7 (2.5-2.9) |
| Female | 2.8 (2.6-2.9) | 2.7 (2.5-2.9) | 2.7 (2.5-2.9) | 2.7 (2.5-2.9) |
| Head circumference (cm) |  |  |  |  |
| Male | 34.0 (33.5-35.0) | 34.0 (34.0-35.0) | 34.0 (34.0-35.0) | 34.0 (33.0-35.0) |
| Female | 34.0 (34.0-35.0) | 34.0 (33.0-35.0) | 34.0 (33.0-35.0) | 34.0 (32.0-35.0) |
| Apgar score<sup>c,d</sup> |  |  |  |  |
| 1 min |  |  |  |  |
| Male | 9 (8-9) | 9 (8-9) | 8 (8-9) | 9 (8-9) |
| Female | 9 (8-9) | 9 (8-9) | 9 (8-9) | 8 (8-9) |
| 5 min |  |  |  |  |
| Male | 9 (9-10) | 10 (9-10) | 9 (9-10) | 10 (9-10) |
| Female | 9 (9-10) | 9 (9-10) | 10 (9-10) | 9 (9-10) |

N, number of individuals; SD, standard deviation; IQR, interquartile range; no., number of events.
<sup>a</sup> Length and Rohrer index was recorded in 157 newborns from non-infected pregnant women.
<sup>b</sup> The Rohrer index is the newborns' weight in grams divided by the cube of the length in centimeters, and babies are considered proportional when values are above 2.5.
<sup>c</sup> Apgar score: 7 – 10, normal; 4 – 6, some breathing assistance might be required; and, < 4, several assistances must be provided.
<sup>d</sup> Apgar score at 1 and 5 minutes was recorded in 153 newborns from non-infected, 112 *P. vivax*, 58 mixed-infected and 52 *P. falciparum* pregnant women.

**Additional file 3: Summary of newborns characteristics of the Prospective Cohort Study according with head circumference.**

**Table 3.** Baseline Characteristics of Newborns at Delivery of Non-Infected and *P. falciparum-infected* Pregnant Women

| Newborns' Characteristics | Median (IQR) |  |  |  |  |
| --- | --- | --- | --- | --- | --- |
|  | Non-Infected<br>(N:M=59, F=79) | NI-SH<br>(N:M=13, F=7) | *Pf*-NHC<br>(N:M=49, F=45) | *Pf*-SH<br>(N:M=15, F=15) | *Pf*-MC<br>(N:M=3, F=5) |
| Weight (g) |  |  |  |  |  |
| Male | 3305.0 (3045.0-3530.0) | 2900.0 (2800.0-3125.0) | 3320.0 (3135.0-3530.0) | 3030.0 (2790.0-3160.0) | 2790.0 (2735.0-4300.0) |
| Female | 3400.0 (3055.0-3690.0) | 3000.0 (2890.0-3240.0) | 3170.0 (2960.0-3390.0) | 2910.0 (2700.0-3100.0) | 2905.0 (2870.0-2910.0) |
| Length (cm)<sup>a</sup> |  |  |  |  |  |
| Male | 49.0 (48.0-50.0) | 48.0 (48.0-49.0) | 50.0 (48.0-51.0) | 49.0 (48.0-50.0) | 49.0 (48.0-54.0) |
| Female | 50.0 (49.0-50.0) | 48.0 (47.0-50.0) | 49.0 (48.0-50.0) | 48.0 (47.0-50.0) | 48.0 (47.0-49.0) |
| Rohrer index<sup>a, b</sup> |  |  |  |  |  |
| Male | 2.8 (2.6-3.0) | 2.6 (2.4-2.7) | 2.7 (2.6-3.0) | 2.5 (2.4-2.7) | 2.5 (2.3-2.7) |
| Female | 2.8 (2.6-2.9) | 2.7 (2.4-3.0) | 2.7 (2.5-2.9) | 2.6 (2.3-2.8) | 2.6 (2.3-2.8) |
| Head circumference (cm) |  |  |  |  |  |
| Male | 35.0 (34.0-36.0) | 33.0 (32.0-33.0) | 35.0 (34.0-35.0) | 32.0 (32.0-33.0) | 32.0 (31.0-32.0) |
| Female | 34.0 (34.0-35.0) | 32.0 (31.0-32.5) | 34.0 (34.0-35.0) | 32.0 (31.0-32.0) | 30.0 (30.0-31.0) |
| Apgar score<sup>c, d</sup> |  |  |  |  |  |
| 1 min |  |  |  |  |  |
| Male | 8.0 (8.0-9.0) | 9.0 (8.5-9.0) | 8.0 (8.0-9.0) | 9.0 (8.0-9.0) | 8.5 (8.0-9.0) |
| Female | 9.0 (8.0-9.0) | 8.0 (6.0-9.0) | 8.0 (8.0-9.0) | 9.0 (8.5-9.0) | 9.0 (9.0-9.0) |
| 5 min |  |  |  |  |  |
| Male | 9.0 (9.0-10.0) | 10.0 (9.0-10.0) | 9.0 (9.0-10.0) | 10.0 (9.0-10.0) | 9.5 (9.0-10.0) |
| Female | 9.0 (9.0-10.0) | 9.0 (9.0-10.0) | 9.0 (9.0-10.0) | 10.0 (9.5-10.0) | 10.0 (10.0-10.0) |

IQR, interquartile range, N, number of newborns; M, male newborns; F, female newborns, NI-SH, Non-Infected small head; *Pf*-NHC, *Plasmodium falciparum*-normal head circumference; *Pf*-SH, *Plasmodium falciparum*-small head; *Pf*-MC, *Plasmodium falciparum*-microcephaly.
<sup>a</sup> Length and Rohrer index were recorded in 58 males from the Non-infected group.
<sup>b</sup> The Rohrer index is the newborns' weight in grams divided by the cube of the length in centimeters, and babies are considered proportional when values are above 2.5.
<sup>c</sup> Apgar score was recorded in 57 males and 77 females from the Non-infected group; in 12 males from the NI-SH group; in 45 males and 39 females from the *Pf*-NHC group; in 14 males and 12 females from the *Pf*-SH group; and, in 2 males from the *Pf*-MC group.
<sup>d</sup> Apgar score: 7 – 10, normal; 4 – 6, some breathing assistance might be required; and, < 4, several assistances must be provided.

**Additional file 4: Summary of serological screening of TORCH, HIV, Syphilis, Dengue, Chikungunya and Zika infections.**

**Table 1.**
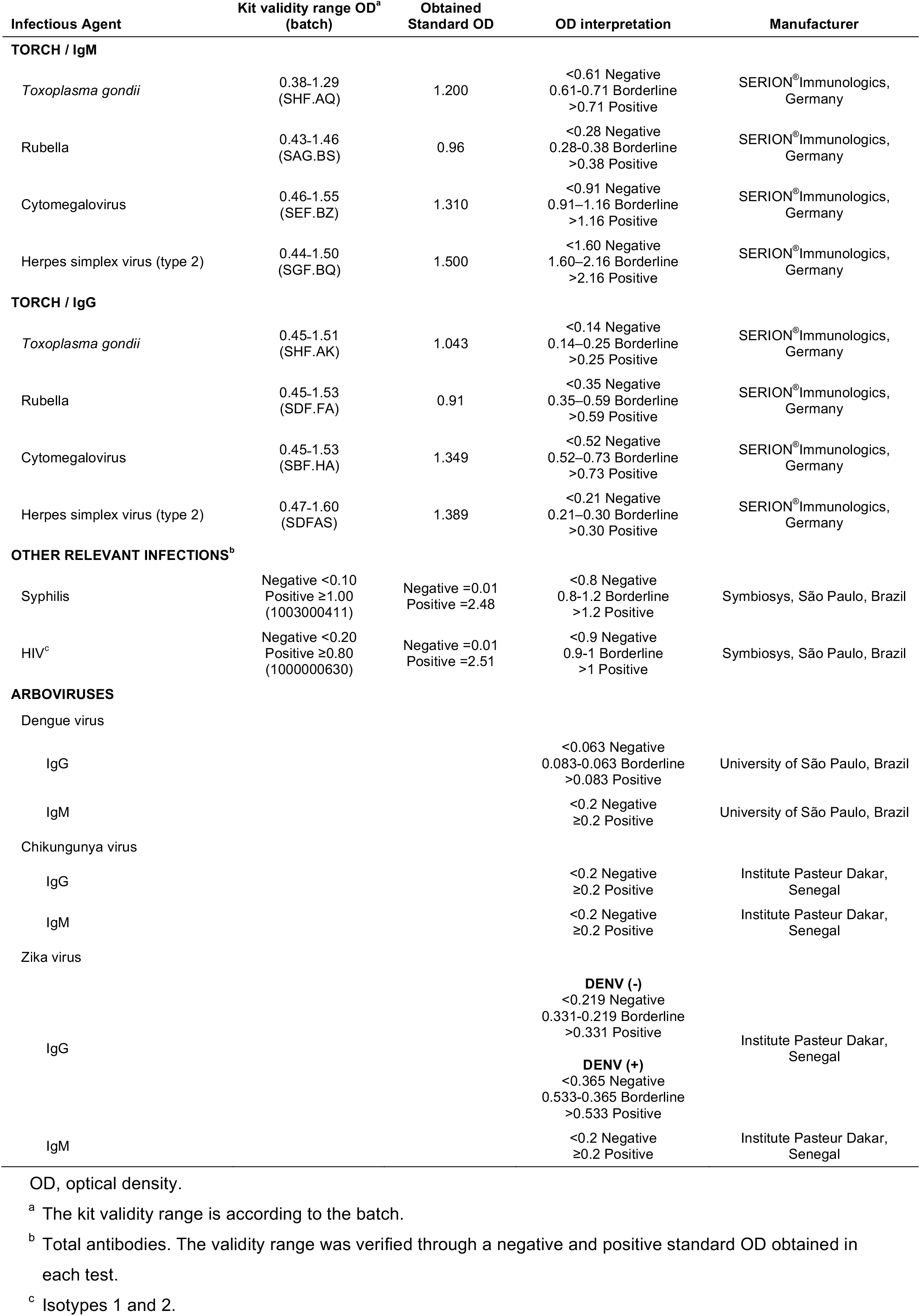
Summary of the serological screening of other infectious agents

**Table 2.**
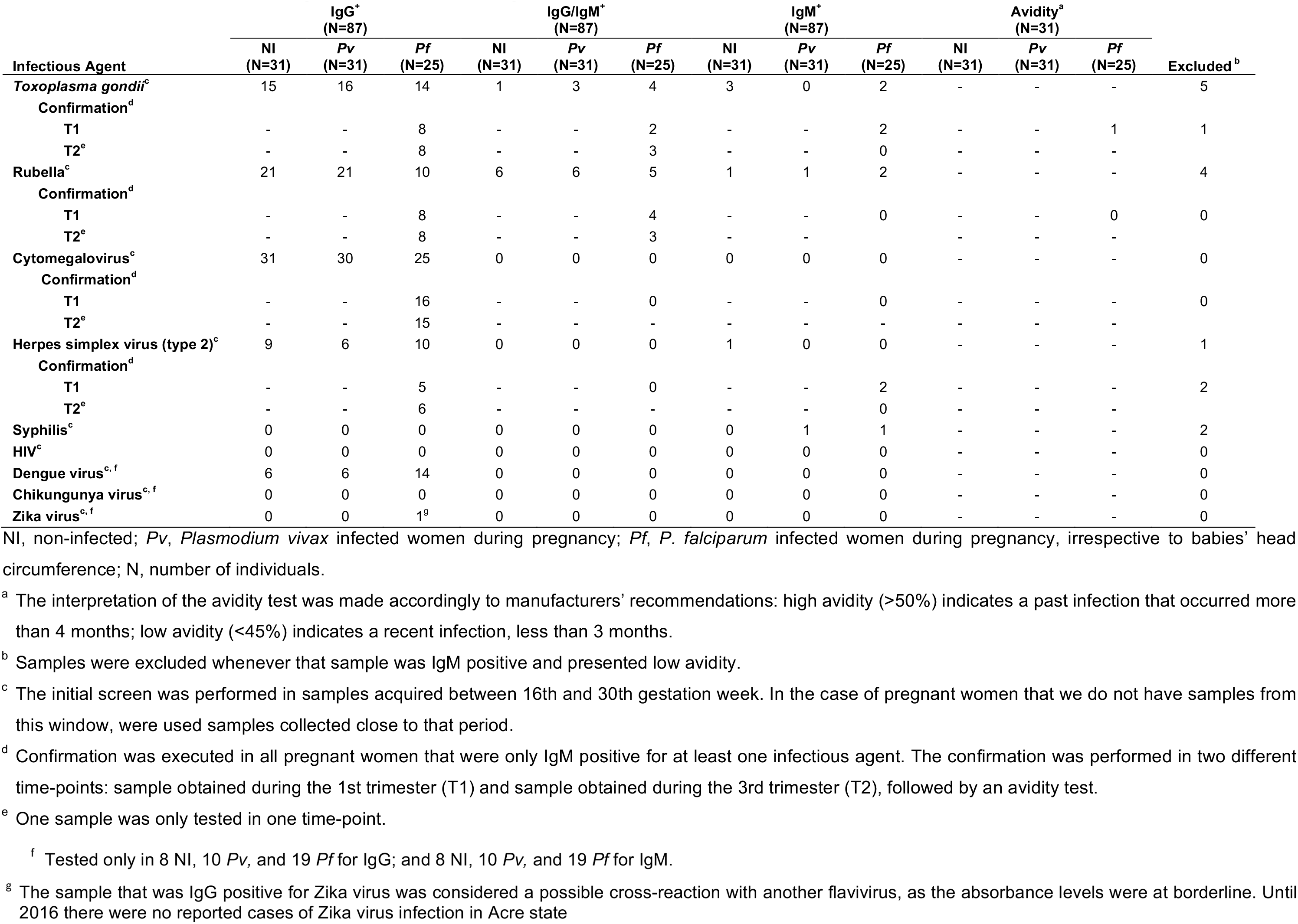
Results of the screening of other infectious agents in mothers of babies with SH in the prospective cohort.

**Additional file 5: Summary of placental parameters evaluation in the Prospective Cohort study according to newborns head circumference.**

**Table 1.** Placental histological parameters and angiogenic factors of non-infected and *P. falciparum-infected* pregnant women

|  | Median (IQR) |  |  |  |  |  |  |  |  |
| --- | --- | --- | --- | --- | --- | --- | --- | --- | --- |
| Characteristics | Non-Infected<br>(N=128) | Non-Infected-SH<br>(N=20) | p-Value<sup>a</sup> | Pf-NHC<br>(N=80) | p-Value<sup>b</sup> | Pf-SH<br>(N=24) | p-Value<sup>c</sup> | Pf-MC<br>(N=7) | p-Value<sup>d</sup> |
| Placental histological parameters |  |  |  |  |  |  |  |  |  |
| Leukocytes infiltrate<sup>e</sup> | 16.0 (9.0-21.0) | 14.0 (10.5-24.0) | 0.905 | 20.0 (13.0-31.0) | 0.001 | 18.0 (14.0-35.0) | 0.073 | 18.0 (17.0-52.0) | 0.083 |
| Monocytes infiltrate<sup>f</sup> | 4.0 (2.0-7.0) | 3.0 (2.0-4.5) | 0.325 | 7.0 (5.0-13.0)<sup>g</sup> | <0.0001 | 9.5 (5.5-15.0)<sup>h</sup> | <0.0001 | 9.0 (6.0-11.0)<sup>i</sup> | 0.018 |
| Fibrin deposition score | 1.9 (1.9-2.4) | 1.9 (1.9-2.8) | 0.446 | 2.4 (1.9-2.8) | <0.0001 | 2.0 (1.9-2.8) | 0.196 | 2.1 (1.9-2.8) | 0.557 |
| Fibrinoid necrosis<sup>j</sup> | 6.0 (3.8-10.2) | 7.2 (4.5-10.3) | 0.707 | 7.1 (4.3-10.0) | 0.425 | 6.8 (4.0-9.8) | 0.898 | 8.1 (2.9-9.4) | 0.937 |
| Proliferation index<sup>k</sup> | 3.6 (2.5-4.9) | 3.6 (2.5-4.4) | 0.281 | 3.8 (2.8-4.8) | 0.613 | 3.7 (2.9-4.2) | 0.716 | 3.0 (2.6-3.5) | 0.208 |
| Villous vascularity<sup>l</sup> | 4.0 (3.6-4.5) | 3.9 (3.4-4.4) | 0.553 | 4.0 (3.4-4.5) | 0.887 | 4.1 (3.6-4.6) | 0.321 | 3.8 (3.6-3.9) | 0.351 |
| Syncytial nuclear aggregates | 13.0 (10.0-17.0) | 14.5 (9.5-16.5) | 0.588 | 13.5 (10.0-19.0) | 0.352 | 17.5 (12.0-24.5)<sup>m, n</sup> | 0.002 | 18.0 (12.0-30.0) | 0.023 |
| Angiogenic factors (ng/mL)<sup>o</sup> |  |  |  |  |  |  |  |  |  |
| ANG-1<sup>p</sup> | 14.9 (10.2-21.1) | 20.8 (14.2-28.8) | 0.024 | 17.7 (10.6-25.9) | 0.047 | 18.1 (11.2-22.3) | 0.275 | 19.4 (10.4-51.7) | 0.468 |
| ANG-2 | 7.0 (3.5-12.8) | 4.6 (3.0-11.5) | 0.465 | 4.2 (1.8-9.1) | 0.007 | 5.5 (2.2-10.0) | 0.088 | 5.1 (0-8.9) | 0.120 |
| ANG-2/ANG-1 ratio | 0.4 (0.2-1.0) | 0.3 (0.1-0.7) | 0.089 | 0.3 (0.1-0.7) | 0.010 | 0.2 (0.1-0.8) | 0.104 | 0.2 (0-0.5) | 0.077 |
| Tie-2 | 13.2 (8.7-18.1) | 16.6 (14.8-21.2) | 0.075 | 16.2 (12.2-21.9) | 0.002 | 15.1 (11.7-21.7) | 0.074 | 18.5 (11.8-22.1) | 0.154 |
| Tie-2/ANG-1 ratio | 0.9 (0.6-1.3) | 0.7 (0.6-1.1) | 0.456 | 0.9 (0.6-1.5) | 0.384 | 0.9 (0.7-1.5) | 0.520 | 0.9 (0.4-2.7) | 0.940 |
| VEGF<sup>p</sup> | 3.2 (1.5-4.6) | 4.3 (2.5-5.4) | 0.032 | 3.7 (2.4-4.5) | 0.097 | 2.9 (1.6-4.7) | 0.834 | 4.8 (4.5-5.7) | 0.009 |
| VEGFR1 | 89.6 (38.9-123.3) | 60.4 (11.5-121.3) | 0.280 | 81.1 (16.4-133.1) | 0.715 | 74.7 (13.0-121.8) | 0.494 | 54.2 (17.2-60.6) | 0.071 |
| VEGFR-2 | 14.2 (12.7-15.8) | 14.7 (13.5-15.6) | 0.406 | 14.4 (13.3-15.7) | 0.359 | 14.3 (12.3-16.2) | 0.868 | 13.6 (12.4-16.1) | 0.611 |
| Leptin (ng/mL)<sup>q</sup> | 33.1 (17.2-47.4) | 22.5 (10.7-37.5) | 0.127 | 21.2 (13.8-42.3) | 0.051 | 19.5 (4.5-37.2) | 0.013 | 16.7 (9.0-26.7) | 0.027 |

N, number of individuals; Non-Infected-SH, non-infected-small head; *Pf*-NHC, *Plasmodium falciparum*-normal head circumference; *Pf*-SH, *Plasmodium falciparum*-small head; *Pf*-MC, *Plasmodium falciparum*-microcephaly; IQR, interquartile range.
<sup>a</sup> Differences between Non-Infected and Non-Infected-SH groups were evaluated using Mann-Whitney rank sum tests.
<sup>b</sup> Differences between Non-Infected and *Pf*-NHC groups were evaluated using Mann-Whitney rank sum tests.
<sup>c</sup> Differences between Non-Infected and *Pf*-SH groups were evaluated using Mann-Whitney rank sum tests.
<sup>d</sup> Differences between Non-Infected and *Pf*-MC groups were evaluated using Mann-Whitney rank sum tests.
<sup>e</sup> Leukocyte infiltrate (CD45+) was recorded in placentas from 126 non-infected, 54 *Pf*-NHC, 17 *Pf*-SH and 5 *Pf*-MC pregnant women.
<sup>f</sup> Monocytes infiltrate (CD68+) was recorded in placentas from 127 non-infected pregnant women.
<sup>g</sup> Statistical difference for the comparison of Non-Infected-SH versus *Pf*-NHC, $p < 0.0001$ .
- <sup>h</sup> Statistical difference for the comparison of Non-Infected-SH versus *Pf*-SH, $p = 0.0005$ . - <sup>i</sup> Statistical difference for the comparison of Non-Infected-SH versus *Pf*-MC, $p = 0.028$ . - <sup>j</sup> Fibrinoid necrosis was recorded in placentas from 78 *Pf*-NHC pregnant women. - <sup>k</sup> Proliferation index was recorded in placentas from 126 non-infected and 77 *Pf*-NHC pregnant women. - <sup>l</sup> Villous vascularity was recorded in placentas from 124 non-infected, 71 *Pf*-NHC, and 23 *Pf*-SH pregnant women. - <sup>m</sup> Statistical difference for the comparison of Non-Infected-SH versus *Pf*-SH groups, $p = 0.050$ . - <sup>n</sup> Statistical difference for the comparison of *Pf*-NHC versus *Pf*-SH groups, $p = 0.023$ . - <sup>o</sup> Angiogenic factors were recorded in placental plasma from 126 non-infected, 18 non-infected-SH and 79 *Pf*-NHC pregnant women. VEGF denotes vascular endothelial growth factor A, VEGFR1 and VEGFR-2 vascular endothelial growth factor A receptor 1 and 2, Ang-1 and 2 angiopoietin-1 and 2. - <sup>p</sup> ANG-1 and VEGF was recorded in placental plasma from 19 non-infected-SH pregnant women. - <sup>q</sup> Leptin was recorded in placental plasma from 126 non-infected, 18 non-infected-SH, 77 *Pf*-NHC and 23 *Pf*-SH pregnant women.

**Table 2.** Inflammatory factors in placental plasma from non-infected and *P. falciparum-infected* women

|  | Median (IQR) |  |  |  |  |  |  |  |  |
| --- | --- | --- | --- | --- | --- | --- | --- | --- | --- |
| Characteristics | Non-Infected<br>(N=126) | Non-Infected-SH<br>(N=19) | *p*-Value<sup>a</sup> | *Pf*-NHC<br>(N=76) | *p*-Value<sup>b</sup> | *Pf*-SH<br>(N=19) | *p*-Value<sup>c</sup> | *Pf*-MC<br>(N=7) | *p*-Value<sup>d</sup> |
| Cytokines (pg/mL)<sup>e</sup> |  |  |  |  |  |  |  |  |  |
| IL1B | 4.8 (4.0-6.0) | 4.7 (3.9-5.5) | 0.548 | 5.8 (4.4-10.7)<sup>f</sup> | 0.0008 | 4.9 (4.2-6.5) | 0.550 | 4.7 (3.9-6.7) | 0.992 |
| IL6 | 76.5 (35.8-146.8) | 54.1 (34.3-121.9) | 0.772 | 114.9 (39.4-207.5) | 0.107 | 132.6 (34.9-187.6) | 0.521 | 40.7 (28.5-157.6) | 0.821 |
| IL8 | 25.5 (15.7-52.2) | 34.2 (12.6-49.9) | 0.716 | 47.2 (23.9-79.0) | 0.0009 | 45.1 (22.1-85.9) | 0.044 | 32.8 (15.0-110.1) | 0.604 |
| IL10 | 4.0 (3.2-4.9) | 3.9 (3.4-5.1) | 1.000 | 6.2 (4.5-10.7)<sup>g</sup> | <0.0001 | 4.1 (3.7-5.1)<sup>h</sup> | 0.222 | 4.1 (4.0-4.4) | 0.503 |
| IL12 | 3.7 (3.0-4.3) | 3.8 (3.3-4.1) | 0.792 | 3.7 (3.0-5.0) | 0.125 | 3.5 (3.2-3.8) | 0.262 | 3.5 (3.4-3.8) | 0.632 |
| TNF | 5.4 (4.5-6.3) | 5.3 (4.2-6.0) | 0.790 | 6.0 (4.7-9.5) | 0.006 | 5.1 (4.2-6.0)<sup>i</sup> | 0.267 | 5.1 (4.2-5.4) | 0.326 |
| Anaphylotoxins (pg/mL)<sup>j</sup> |  |  |  |  |  |  |  |  |  |
| C3a | 4.5 (3.2-6.6) | 4.6 (3.4-6.5) | 0.981 | 2.1 (0-5.3)<sup>k</sup> | <0.0001 | 3.0 (0-5.5) | 0.014 | 3.6 (0-6.5) | 0.406 |
| C4a | 37.1 (22.4-54.1) | 30.2 (18.5-39.0) | 0.108 | 27.2 (16.6-44.8) | 0.017 | 24.9 (16.7-49.9) | 0.100 | 29.3 (16.7-52.5) | 0.397 |
| C5a | 1164.0 (835.2-1552.4) | 879.6 (617.4-1065.9) | 0.002 | 1576.9 (1086.3-2384.2)<sup>l</sup> | <0.0001 | 1292.3 (830.6-1618.2)<sup>m</sup> | 0.512 | 1326.4 (830.6-1510.3)<sup>n</sup> | 1.000 |

Non-Infected-SH- Non-Infected-small head; *Pf*-NHC, *Plasmodium falciparum*-normal head circumference; *Pf*-SH, *Plasmodium falciparum*-small head; *Pf*-MC, *Plasmodium falciparum*-microcephaly; N, number of individuals; IQR, interquartile range.
<sup>a</sup> Differences between Non-Infected and Non-Infected-SH groups were evaluated using Mann-Whitney rank sum tests.
<sup>b</sup> Differences between Non-Infected and *Pf*-NHC groups were evaluated using Mann-Whitney rank sum tests.
<sup>c</sup> Differences between Non-Infected and *Pf*-SH groups were evaluated using Mann-Whitney rank sum tests.
<sup>d</sup> Differences between Non-Infected and *Pf*-MC groups were evaluated using Mann-Whitney rank sum tests.
<sup>e</sup> IL1B denotes interleukin-1 beta; IL6, interleukin-6; IL8, interleukin-8; IL10, interleukin-10; IL12, interleukin-12; and TNF, tumor necrosis factor $\alpha$ .
<sup>f</sup> Statistical difference for the comparison of Non-Infected-SH versus *Pf*-NHC groups, $p = 0.015$ .
<sup>g</sup> Statistical difference for the comparison of Non-Infected-SH versus *Pf*-NHC groups, $p < 0.0001$ .
<sup>h</sup> Statistical difference for the comparison of *Pf*-NHC versus *Pf*-SH groups, $p = 0.003$ .

**Additional file 6: Summary of maternal and newborns characteristics of the Retrospective Cohort Study (RCS).**

**Table 1.** Baseline characteristics of mothers and newborns

| Characteristics | Non-Infected<br>(N=3650) | *P. vivax*<br>(N=129) | Mixed<br>(N=21) | *P. falciparum*<br>(N=82) |
| --- | --- | --- | --- | --- |
| **Mothers** |  |  |  |  |
| Maternal age (years), mean (SD)<sup>a</sup> | 24.1 (6.4) | 22.7 (6.1) | 23.4 (6.4) | 24.1 (6.6) |
| Gravidity, no. (%)<sup>b</sup> |  |  |  |  |
| Primigravida | 1286 (36.4) | 46 (36.2) | 6 (28.6) | 22 (28.2) |
| Multigravida | 2247 (63.6) | 81 (63.8) | 15 (71.4) | 56 (71.8) |
| Gestational age at delivery (weeks) |  |  |  |  |
| Mean (SD) | 39.3 (1.1) | 39.0 (1.2) | 39.2 (1.4) | 39.0 (1.1) |
| Median (IQR) | 39.0 (38.0-40.0) | 39.0 (38.0-40.0) | 39.0 (38.0-40.0) | 39.0 (38.0-40.0) |
| C-section, no. (%) | 1283 (35.2) | 43 (33.3) | 7 (33.3) | 22 (26.8) |
| Antenatal care visits, mean (SD)<sup>c</sup> | 5.9 (2.5) | 5.8 (2.4) | 5.6 (2.4) | 5.5 (2.8) |
| **Newborns** |  |  |  |  |
| Male newborns, no. (%) | 1921 (52.6) | 67 (51.9) | 10 (47.6) | 47 (57.3) |
| Weight (g), median (IQR) |  |  |  |  |
| Male | 3325.0 (3065.0-3650.0) | 3360.0 (3005.0-3600.0) | 3242.5 (3165.0-3750.0) | 3180.0 (2960.0-3470.0) |
| Female | 3200.0 (2965.0-3480.0) | 3150.0 (2910.0-3410.0) | 3035.0 (2700.0-3340.0) | 3100.0 (2860.0-3380.0) |
| Length (cm), median (IQR)<sup>d</sup> |  |  |  |  |
| Male | 49.0 (48.0-51.0) | 49.0 (48.0-50.0) | 49.0 (48.0-51.0) | 49.0 (48.0-50.0) |
| Female | 49.0 (48.0-50.0) | 48.0 (48.0-49.0) | 49.0 (48.0-50.0) | 48.0 (47.0-49.0) |
| Rohrer index, median (IQR)<sup>d, e</sup> |  |  |  |  |
| Male | 2.8 (2.6-3.0) | 2.8 (2.6-3.0) | 2.8 (2.5-3.0) | 2.8 (2.6-3.0) |
| Female | 2.8 (2.6-3.0) | 2.8 (2.6-2.9) | 2.4 (2.3-2.7) | 2.7 (2.5-3.0) |
| Head circumference (cm), median (IQR) |  |  |  |  |
| Male | 34.0 (33.0-35.0) | 34.0 (33.0-35.0) | 34.5 (33.0-36.0) | 34.0 (33.0-35.0) |
| Female | 34.0 (33.0-35.0) | 34.0 (33.0-35.0) | 34.0 (32.0-35.0) | 33.0 (32.0-35.0) |
| Apgar score, median (IQR)<sup>g, h</sup> |  |  |  |  |
| 1 min |  |  |  |  |
| Male | 9 (8-9) | 9 (8-9) | 8 (8-9) | 9 (8-9) |
| Female | 9 (8-9) | 9 (8-9) | 9 (8-9) | 8 (8-9) |
| 5 min |  |  |  |  |
| Male | 10 (9-10) | 10 (9-10) | 9 (9-10) | 10 (9-10) |
| Female | 10 (9-10) | 10 (9-10) | 10 (9-10) | 10 (9-10) |

N, number of individuals; SD, standard deviation; IQR, interquartile range; no., number of events.
- <sup>a</sup> Maternal age was recorded in 3372 non-infected and 126 *P. vivax*-infected pregnant women. - <sup>b</sup> Gravidity was recorded in 3533 non-infected, 127 *P. vivax*, and 78 *P. falciparum*-infected pregnant women. - <sup>c</sup> The number of antenatal care visits was recorded in 3413 non-infected, 125 *P. vivax*, 20 mixed-infected and 79 *P. falciparum* pregnant women. - <sup>d</sup> Length and Rohrer index was recorded in 3635 newborns from non-infected pregnant women. - <sup>e</sup> The Rohrer index is the newborns' weight in grams divided by the cube of the length in centimeters, and babies are considered proportional when values are above 2.5. - <sup>f</sup> Chest circumference was recorded in 3647 newborns from non-infected pregnant women. - <sup>g</sup> Apgar score: 7 – 10, normal; 4 – 6, some breathing assistance might be required; and, < 4, several assistances must be provided. - <sup>h</sup> Apgar score at 1 and 5 minutes was recorded in 3628 newborns from non-infected and 81 *P. falciparum*-infected pregnant women

